# Materials on Plant Leaf Surfaces are Deliquescent in a Range of Environments

**DOI:** 10.1101/2021.11.22.469518

**Authors:** E. C. Tredenick, H. Stuart-Williams, T. G. Enge

## Abstract

Materials on plant leaf surfaces that attract water impact penetration of foliar-applied agrochemicals, foliar water uptake, gas exchange and stomatal density. Few studies are available on the nature of these substances, and we quantify the hygroscopicity of these materials. Water vapor sorption experiments on twelve leaf washes of sample leaves were conducted and analyzed with inductively coupled plasma-optical emission spectroscopy and X-ray diffraction. All leaf surface materials studied were hygroscopic. Oils were found on the surface of the *Eucalyptus* studied. For mangroves that excrete salt to the leaf surfaces, significant sorption occurred at high humidity of a total of 316 mg (∼ 0.3 mL) over 6–10 leaves, and fitted a Guggenheim, Anderson, and de Böer sorption isotherm. Materials on the plant leaf surface can deliquesce and form an aqueous solution in a variety of environments where plants grow, including glasshouses and by the ocean, which is an important factor when considering plant-atmosphere relations.

## 1. Introduction

The deposition of aerosols onto plant leaves is a common occurrence in the environment, yet its ecophysiological impacts are poorly understood. The aerosols are highly variable in composition and size (1 nm to 100 *μ*m [37]). The impact of aerosols deposited onto plant leaves requires further attention and few studies are available on the implications of deposited aerosols on plants in regard to plant-atmosphere relations, plant physiology and micrometeorology. We aim to test if hygroscopic particles on plant leaf surfaces are present in a range of environments and attempt to characterize them. Hygroscopic particles are contained in aerosols and may be deposited onto plant leaf surfaces, impacting many factors, both in experimental settings and field work. The deposition of aerosols of calcium has increased in the western USA, due to mineral aerosols from dust storms, increased human activity up-wind, increased aridity and wind transport [5]. Leaf surface wetness can increase trace gas deposition and provide a trap for easily soluble compounds [34, 42]. Ionic substances are often present on plant leaves (especially in saline environments, such as mangroves [17]), in atmospheric particles and in sprinkler irrigation [25, 33]. Hygroscopic materials can reduce the surface tension of droplets [22] and may allow stomatal penetration [11, 14, 23]. Gas exchange experiments, stomatal apertures and distribution [14, 28], foliar water uptake [17], and foliar-applied agrochemical penetration [47] may be impacted by hygroscopic materials. In the context of foliar-applied agro-chemical spray penetration, the effect of additional hygroscopic materials on the surface is significant, including changing the point of deliquescence of the applied salt [16, 25], total droplet evaporation time, droplet contact angle and area, and total amount of chemical penetrated [46–49].

The average thickness of a liquid layer present on a leaf surface due to hygroscopic particles is estimated to be approximately 1 *μ*m [11]. Particulate matter present on leaf surfaces can reach a similar mass to that of leaf waxes; wax mass is around 50 *μ*g cm^*−*2^ [11, 41]. The electrical conductance, related to surface wetness, for a leaf and artificial leaf in a field, is similar at night (leaf stomata closed), while during the day (leaf stomata open) the leaf is more conductive. Stomata opening during the day play an important role in controlling leaf moisture [10, 11].

Foliar water uptake may be important for the plant during drought. A large proportion of plant species studied have been found to have a capacity for foliar water uptake, totaling 124 species [18]. Water may be present on the leaf surface from a variety of sources including rain, dew and high humidity. Fog suppresses water loss from leaves, for example ameliorating daily water stress in a coastal redwood. A diurnal rhythm is present and older, well watered leaves take up the most water [8]. Without fog, species with high foliar water uptake are more likely to lose turgor during seasonal droughts [24].

A variety of mangrove species have glands in their leaves that excrete salts to the leaf surface, as shown for a mangrove grown in a growth cabinet (Fig. 1). These salt crystals are large and visible to the naked eye. Mangroves are known to rely on non-saline water to maintain productivity and foliar water uptake may be key. Three species of mangroves growing in arid and humid environments have been shown to have a contribution from foliar water uptake of 32% in *Avicennia germinans*, 26% in *Laguncularia racemosa* and 16% in *Rhizophora mangle*, and of these, *Avicennia germinans* excretes salts onto the leaf surface [31]. For these plants to achieve full leaf hydration, they require the input of water from additional sources other than root water, such as atmospheric water [38]. Within the same species, uptake was comparable across field and controlled environments, suggesting that uptake is not a plastic arid-zone adaptation but may be used as a supplemental water balance strategy in humid and arid neotropical mangroves [31]. Leaf water potential and reverse sap flow rate increase above the point of deliquescence (POD) (75%RH for sodium chloride, NaCl) [17], indicating that surface salts are an important consideration in foliar water uptake for mangroves. Plants living in halophilic environments, such as saltbush (*Atriplex halimus*) [42] excrete salt onto the leaf surface. Species with hygroscopic salts on the leaf surface have leaves that will stay wet longer and trap aerosols. Species with more NaCl on their leaves have very limited diversity of bacteria and fungi on the leaf surface, for example on saltbush (*Atriplex halimus*) and mangroves (*Avicennia germinans and Laguncularia racemosa*) [26, 42], indicating that salt on plant leaves may be a defense mechanism. Thus the presence of salts on the leaves may yield multiple benefits, including improved water relations and bacterial control.

**Figure 1:**
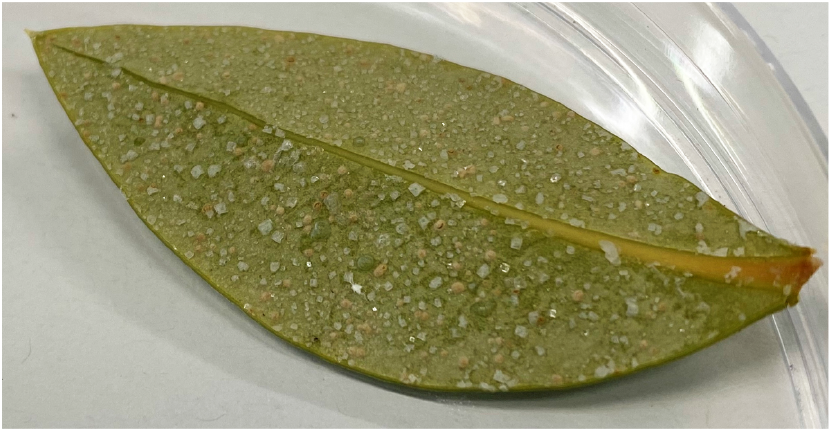
Naturally occurring salts on the freshly cut abaxial leaf surface of ‘Cabinet mangrove’ (grey/white mangrove *Avicennia marina*). The salts are present on the leaf due to glands that excrete salt and the leaf is visibly dry.

Ionic substances can sorb water from the air, due to their strong attractive forces for highly polar water molecules. The ability to attract (adsorb or absorb) water molecules from the air is termed hygroscopicity. The point, in terms of relative humidity, at which a hygroscopic material can sorb enough water to dissolve and form an aqueous solution is called the point of deliquescence (POD) or deliquescent relative humidity (DRH). Most deliquescent materials are ionic salts and their POD can vary significantly at room temperature, from 32%RH for CaCl_2_ [35], 75%RH for NaCl, to 97%RH for K_2_SO_4_ [43]. If the relative humidity is above the POD, solid crystals will sorb moisture from the air until the salt dissolves and remains in solution [21, 39]. The solution will continue to sorb water from the air until an equilibrium is reached between the vapor pressure of the solution and the air. For example, if the relative humidity of the air is 50%RH and is above the POD of CaCl_2_ (32%RH), salt crystals or salt solution will attract water. If the relative humidity is below the POD, say 10%RH, the solution will continue to evaporate and eventually form crystals. Additional to relative humidity, the rate of sorption by hygroscopic material depends on temperature, the surface area of the salt exposed to the air and the wind or air circulating over it [21]. NaCl crystals are seen under fluctuating relative humidities around the POD of NaCl on astomatous isolated tomato fruit cuticles, and pools of water form [9]. The particular point of deliquescence of many ionic salts is well defined, though there is less known about the sorption of salt mixtures, and salt and oil mixtures. Plant matter sorption is significantly less than hygroscopic salts, where sorption at high humidity is 30% in cellulose, 49% in polar polysaccharides isolated from cuticles and 21% in clays (smectite), while CaCl_2_ can sorb 1400% [3, 20, 21, 36].

Heterogeneous stomatal pore area or patchy stomatal conductance may have substantial implications for photosynthetic efficiency. Plants grown in filtered or unfiltered air were compared and aerosols deposited onto the leaf from unfiltered air suppressed the heterogeneity of stomatal pore opening and response to vapor pressure deficit [27] while increasing the minimum epidermal conductance [14]; a key factor of drought tolerance in plants [13]. Hygroscopic aerosols may contribute to the formation of a thin aqueous film across the leaf surface that can connect stomata to each other and the leaf interior [27, 28]. Salts artificially sprayed onto leaves increased the minimum epidermal conductance [13], and the electrical leaf conductance related to leaf surface wetness [12].

We aim to determine whether hygroscopic materials exist on plant leaves, across a wide range of locations where plants grow. We focus on any substance present on the plant leaf surface (*in situ*), which is easily washed off and dissolved in solution, and remains in solution after centrifugation. These substances may originate from deposited aerosols, sea spray, impurities in rain, soils and salts, or inside the leaf due to leaching or excreted salts. We aim to determine the materials’ compositions; how hygroscopic they are over a large range of relative humidities, and whether they visibly deliquesce and form an aqueous solution. Vapor sorption methods are used to analyze the leaf material sorption, while the leaf wash material composition are analyzed using powder X-ray diffraction and inductively coupled plasma-optical emission spectroscopy (ICP-OES). We compare the leaf wash sorption to control sorption experiments conducted alongside.

## 2. Methods

### 2.1. Leaf Wash Sample Preparation

Plant samples were taken at a range of distances from the ocean, with locations and species described in Table 1, and total dry weights in Table 4. These locations included by the ocean, river, farm, lake, indoor, town and inside glasshouses and growth cabinets. The leaves were collected from the same individual plant at each location, and from the same unshaded part of the plant. All samples were collected after at least two weeks without rain and were from mature plants, unless otherwise stated. No artificial spray applications or pre-treatments were used during the experiment.

**Table 1:** Sample reference, plant species and location of leaf wash samples. 6–10 leaves were collected with similar leaf area. Mature plants were collected 2 weeks after rain unless otherwise specified. Locations are in the Australian Capital Territory (ACT) or New South Wales (NSW), Australia. Collection dates between February and July 2020. Growth cabinets were contained in a large air conditioned indoor facility with 210 micron mesh air filtration, rated to P2 standards for genetically modified organisms.

| Reference | Common name | Scientific name | Location | Notes |
| --- | --- | --- | --- | --- |
| Brackish man-grove | river mangrove | *Aegiceras corniculatum* | Currowan, NSW, Aust | 10 km to ocean, edge of river, submerged leaves at certain tides, excretes salt |
| Cabinet mangrove | grey/white mangrove | *Avicennia marina* | growth cabinet | excretes salt, not watered on leaves, 30°C day 20°C night, 60%RH day 70%RH night, 8 months old |
| Glasshouse chilli | birds eye or Thai chilli | *Capsicum annuum* | glasshouse | not watered on leaves, 28°C day 20°C night, around 2 years old |
| Ocean common reed | common reed | *Phragmites australis* | Surf Beach, NSW, Aust | 20m from beach, unlikely to be submerged |
| Cabinet setaria | setaria green foxtail | *Setaria viridis* | growth cabinet | mature, 30°C day 25°C night, 60%RH day 70%RH night, not watered on leaves |
| Brackish euc | eucalyptus grey iron-bark | *Eucalyptus paniculata* | same as brackish mangrove | edge of river, less likely to be submerged |
| Indoor peace lily | white peace lily | *Spathiphyllum cochlearispathum* | indoors | not watered on leaves |
| Cabinet barley | barley | *Hordeum vulgare* | growth cabinet | as setaria |
| Town euc rain | eucalyptus torelliana | *Cadaghi corymbia torelliana* | Acton, ACT, Aust | 6 hours after rain, rained for several days, leaves dry after rain, 200kms inland from coast |
| Town euc no rain | eucalyptus torelliana | *Cadaghi corymbia torelliana* | Acton, ACT, Aust | same plant as Town euc rain |
| Lake euc | long-leaved box eucalyptus | *Eucalyptus goniocalyx* | Acton, ACT, Aust | 20m from lake |
| Euc farm | inland scribbly gum eucalyptus | *Eucalyptus rossii* | Monga, NSW, Aust | Near farms, 10m from dirt road, 100m from highway, halfway between town and ocean |

Leaves of around the same area (6–10 leaves washed in total, with a total leaf area including the 2 washed leaf surfaces of approximately 430 cm^2^), enough to fill a glass container (125 mL) with a screw cap lid, were collected intact on their branch (so a small area of petiole and stem was also washed) and removed carefully from the plant. The leaves were placed in the bottle with 60 mL of de-ionized water, immediately after collection from the plant and shaken lightly for 20 seconds [14]. Then the leaves were quickly removed from the wash solution and discarded. The glass bottle was thoroughly cleaned beforehand, with detergent, then rinsed with de-ionized water and ethanol. The wash was centrifuged at 2,000 g for 15 minutes (Orbital 420, Clements, Australia). Centrifuging was repeated once if particles remained suspended. We note that the wash was not filtered as we wanted to investigate small particles. For each plant species and location, a total of two sample sets were collected from one individual plant, 120 mL in total; one for the vapor experiment and one for the ICP-OES experiment. The ICP-OES sample had the leaf matter removed and discarded, and was then centrifuged and dried from 60 mL to 5 mL.

For the vapor experiment to find the initial dry weights of the vessel, 3 empty 1.5 mL polypropylene Eppendorf tubes were prepared by drying them in the oven and then weighed to determine the vessel dry weights, *dw*_*v*_. For the leaf washes, each 60 mL sample was divided equally into 3. Samples were dried in an oven for 3 days at 50°C. The wash was dried down to a smaller volume, then placed in the dry Eppendorf tubes (the same 3 from the dry weight measurement) and dried further to a solid. The total dry weight of the sample and vessel was recorded, *dw*_*v*+*s*_. Sample mass was determined for all dry weight and sorption experiments using a Mettler AT21 Comparator, Mettler Toledo, Italy, d = 1 *μ*g, max = 22 g.

### 2.2. Static Vapor Gravimetric Sorption Experiment - Leaf Washes

The static vapor gravimetric sorption technique [1] was used to determine the sorption of water by the samples. Wash samples start out oven dry, then are successively placed in environments of increasing relative humidity to test their adsorption properties. To create a specific relative humidity, a range of salt solutions were used, being MgCl_2_, Ca(NO_3_)_2_, NaCl, KCl, KNO_3_, K_2_SO_4_, with the equilibrium RH being 33.4, 55.5, 75, 84, 93 and 97%RH, respectively. To create these saturated salt solutions, 50 mL of de-ionized water was placed in a large petri dish, next to the leaf sample with the cap open, inside an air-tight larger glass vessel. The larger vessel was then stored in a temperature controlled environment for 3 days (21.9 ± 0.4°C). At the end of 3 days, samples were quickly taken out of the large vessel, re-capped and weighed in their Eppendorf tubes to provide the wet weight of the sample, *ww*_*v*+*s*_, at that particular relative humidity. It was noted if liquid was clearly visible over the material in the Eppendorf tube without the need for special equipment. Triplicates were compared and averaged. Error bars were produced for the standard error of the mean. The polypropylene Eppendorf tubes can also sorb moisture, so the wet weight of the tubes was approximated from 6 empty tubes exposed to each humidity.

### 2.3. Salt and Oil Controls - Water Sorption Experiment

We compare the sorption of the leaf washes for similar weights to controls based on salts and oils including mixtures. Salts and oils were weighed as shown in Table A6 and added to 1mL of de-ionized water. The samples were then mixed and dissolved where possible, and dried in the oven at 50°C for 3 days. The same procedure as Section 2.2 was conducted for measuring water sorption. The mangrove nutrient (that the ‘Cabinet mangrove’ grew in a solution of), *Eucalyptus* oil (100% pure) and tea tree oil (100% pure, *Melaleuca alternifolia*, Australia) were purchased. The mangrove nutrient comprises mainly NaCl, along with 1295 ppm Mg, 430 ppm Ca and 390 ppm K.

### 2.4. X-ray Diffraction

Powder X-ray diffraction (XRD) analysis was carried out with a Malvern Panalytical Empyrean Series 3 diffractometer, with Bragg-Brentano^HD^ divergent beam optic and a PIXcel^3D^ detector (1D scanning mode, 3.347° active length), using Co*Kα* radiation. Two samples were analyzed with the leaf wash method (Table 2). Samples were analyzed with a broad beam (long-fine focus) over a range of 4–85° 2*θ*, with a step with of 0.0131303° and scan speeds ranging from 298–2,598 seconds per step depending on sample requirements. Samples were rotated horizontally to increase sampling size. Two other methods were utilized with 6 plant species; by directly studying the leaf, and its scrapings. Samples were obtained by scraping materials off the leaves (Figs. A14, A15) with a scalpel and washing any residual sample into an agate mortar with ethanol, then grinding the material with an agate pestle by hand as finely as possible. The sample was then deposited with a Pasteur pipette onto a low-background sample holder (made of Si or quartz), dried and presented to the X-ray beam without the leaf substrate. The only possible contaminant of samples prepared this way is small amounts of epicuticular wax from the plant (paraffin). Leaf wash samples (see Section 2.1, Table 2 and Figs. A12, A13) were also prepared on such low background holders. The wash samples proved somewhat sub-optimal for XRD analysis because salts readily dissolve and precipitated as potentially different compounds. The suitability of each sample for yielding instructive powder XRD data varied, depending on the sample preparation (leaf wash, leaf scraping or direct analysis of the leaf surface), the amount of material on the leaf surface, the crystal sizes (1–10 *μ*m ideal), and leaf shape (flat better than curled for direct leaf analysis), requiring individual evaluation of analytical conditions and methods for each sample. Phase identification was carried out with the software DiffracPlus Eva 10 [6] and ICDD PDF-2 database [32], and quantification with Siroquant V4 [44].

**Table 2:** XRD results for the two leaf wash samples that were expected to have the most and least amounts of materials present on the surface; ‘Brackish mangrove’ and ‘Town euc rain’. An ‘x’ in the table indicates that significant amounts were present, ‘trace’ indicates presence in small quantities. The XRD profiles are shown in Figs. A12 and A13.

| Minerals | Chemical composition | Brackish mangrove | Town euc rain |
| --- | --- | --- | --- |
| quartz | SiO<sub>2</sub> |  | x |
| plagioclase | (Ca,Na)<sub>1-2</sub>(Si,Al)<sub>2-3</sub>O<sub>8</sub> |  | x |
| K-feldspar | KAlSi<sub>3</sub>O<sub>8</sub> |  | trace |
| illite/muscovite | KAlSi<sub>3</sub>O<sub>10</sub>(OH)<sub>2</sub> |  | x |
| kaolinite | Al<sub>2</sub>Si<sub>2</sub>O<sub>5</sub>(OH)<sub>4</sub> | x | trace |
| 2:1 clay - chlorite, vermiculite or smectite |  |  | x |
| boehmite | AlOOH |  | x |
| halite | NaCl | x |  |
| sylvite | KCl | x | trace |
| talc | Mg<sub>3</sub>(OH)<sub>2</sub>Si<sub>4</sub>O<sub>10</sub> |  | x |
| unidentified phase, possibly quartz |  | x |  |

### 2.5. ICP-OES - Inductively Coupled Plasma-optical Emission Spectroscopy

The quantification of 70 elements was carried out using an Agilent 5110 ICP-OES (Agilent Technologies, Australia), operating in Synchronous Vertical Dual View (SVDV) mode, allowing for the simultaneous detection of axial and radial emission signals. Only concentrations for Al, B, Ca, Cu, Fe, K, Mg, Na, P, Si, Sr and Zn were detected above the method detection limit of 0.1 *μ*g*/*g. A double pass cyclonic spray chamber, a SeaSpray nebuliser, and a 2.4 mm quartz injector were used as the introduction system. Operating parameters for the ICP-OES analysis are tabulated in Table A8. All dilutions and sample preparation of samples for ICP-OES measurement were performed using ultrapure water (MilliQ, Merck), as well as sub-boiling distilled HNO_3_. A custom multi-element calibration solution for the elements of interest was prepared from single element standard solutions (Inorganic Ventures) and diluted to concentrations ranging from 0.1 to 20 *μ*g*/*mL. All samples were diluted and acidified to fall within the calibration curve and repeat analyses were carried out with multiple dilution steps for samples that initially exceeded the calibration range. Blank contribution was monitored by acidifying and analyzing the de-ionized water used for the wash. The presence of chloride (Cl^−^) and sulphate (SO_4_^2−^) was tested, as described in Section A1.

### 2.6. Moisture Gain Calculations

To find the total weight and percentage increase in moisture gained for each relative humidity step for the sample, we consider the dry weight and the blank vessel water sorption. We first consider

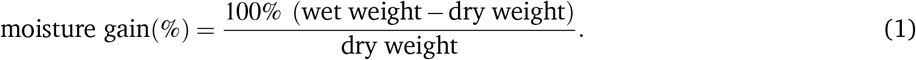

We formulate Δ*w* (%), the weight increase of moisture adsorbed above the dry weight of the sample (sample being the salt taken from the leaf surface) scaled with the blank, as follows:

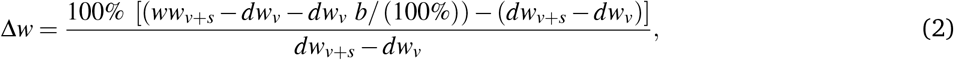

where *ww*_*v*+*s*_ (mg) is the total wet weight of the vessel and sample together at a given RH, *dw*_*v*_ (mg) is the empty vessel/ Eppendorf tube dry weight, *b* (%) is the blank percentage moisture weight gain at a given RH and *dw*_*v*+*s*_ (mg) is the total vessel and sample dry weight. Simplifying, the final equation for Δ*w* becomes:

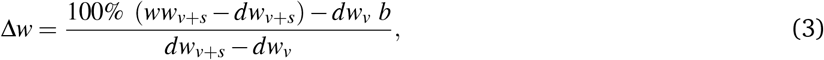

and Total mg, the total weight of the wet sample including the sample dry weight is:

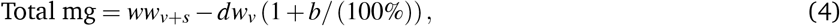

The parameters are direct weight measurements, except Δ*w*, Total mg, and *b*, which are calculated. Dry weights were determined following oven drying. As Eppendorf tubes vary in weight, each tube is weighed dry and with the

sample. To correct for water sorption by the polypropylene, 6 blank Eppendorf tubes were weighed with successive humidity steps, and averaged, to find *b* using Equation (1), producing sorption between 0.033% to 0.23%, at 33%RH to 97%RH.

## 3. Results

To investigate what materials are on the surface of plant leaves, we use a range of techniques, with the plant species and locations described in Table 1. We use the word “materials” due to their highly mixed nature. The total dry weights of the triplicates after centrifugation and drying are shown in Table 4, and indicate that both mangroves have large amount of materials present, 40 and 41 mg (when appropriately scaled), compared to the other samples (3 to 8 mg). The total weight of dry material is relevant in this study, as increasing the hygroscopic material also increases the total sorption weight, and our main interest is the total amount of water that could accumulate on a leaf surface. To investigate the types of materials present on the leaf, our initial analysis was performed using leaf washes with the XRD, as shown in Table 2 and Figs. A12 and A13. ‘Brackish mangrove’ contained significant amounts of ionic hygroscopic compounds such as NaCl and KCl, along with other minerals, while ‘Town euc rain’ contained a range of materials but only trace amounts of hygroscopic KCl. These samples were dissolved in deionized water and represent the compounds on the leaf but may not represent the original *in situ* configuration of the elements, for example NaCl and K_2_SO_4_ might become Na_2_SO_4_ and KCl, having different PODs. Tests were carried out to determine what the original compounds were on the leaf and results were similar to those in Table 2, as shown in Figs. A14 and A15. Scrapings from the leaves gave better results than viewing the leaf directly and significant portions of wax were also present on the leaves but not quantified. XRD is unable to detect and analyze materials without a repetitive or crystalline structure that occur in very small amounts (lower detection limits for XRD range from about 0.1 to 2 wt.% depending on compound). Therefore, ICP-OES analyses were carried out to examine the elemental composition of the samples, including trace elements.

The ICP-OES results are shown in Table 3 in *μ*g*/*g (ppm (parts per million)) and grey shading indicates relatively higher concentrations. The results indicate that the samples having significant amounts of hygroscopic salts are the ‘Brackish mangrove’, ‘Cabinet mangrove’, ‘Brackish euc’, ‘Glasshouse chilli’ and perhaps the ‘Euc farm’ and ‘Ocean common reed’, based on the determined concentration of Ca, Mg and Na. The ‘Brackish mangrove’ and ‘Cabinet mangrove’ have similarly large amounts of most elements. The presence of Cl^−^ was tested separately, indicating ‘Brackish mangrove’, ‘Cabinet mangrove’ and ‘Brackish euc’ contained significant amounts of Cl^−^, consistent with the calculated Na values. The silver nitrate test changed all 5 Eucalyptus samples, containing Eucalyptus oil, brown as seen in Fig. A6. ‘Town euc no rain’ shows the darkest color, indicating the largest amount of oil. Testing for SO_4_^2−^ was also carried out on these samples, with no positive indications.

**Table 3:** ICP-OES results in *μ*g/g (ppm) of similar leaf area for all leaf wash samples studied. Grey shading indicates comparatively higher concentrations. The silver nitrate test for Cl^−^ is described in Section A1. A test for SO_4_^2 –^ was also carried out, with no positive indications.

| Sample | Al | B | Ca | Cu | Fe | K | Mg | Mn | Na | P | Si | Sr | Zn | Cl |
| --- | --- | --- | --- | --- | --- | --- | --- | --- | --- | --- | --- | --- | --- | --- |
| Brackish mangrove | 0.06 | 0.11 | 7.24 | 0.00 | 0.00 | 29.31 | 6.55 | 0.11 | 148.03 | 0.19 | 0.19 | 0.07 | 0.03 | x |
| Cabinet mangrove | 0.03 | 0.16 | 9.78 | 0.06 | 0.01 | 30.20 | 4.46 | 0.04 | 199.20 | 0.15 | 0.23 | 0.03 | 0.09 | x |
| Glasshouse chilli | 0.02 | 0.08 | 6.07 | 0.09 | 0.01 | 4.14 | 3.63 | 0.05 | 0.98 | 0.62 | 0.22 | 0.01 | 0.06 |  |
| Ocean common reed | 0.03 | 0.07 | 3.20 | 0.04 | 0.01 | 5.88 | 3.09 | 0.09 | 3.02 | 0.05 | 0.27 | 0.05 | 0.17 |  |
| Cabinet setaria | 0.01 | 0.09 | 3.91 | 0.06 | 0.00 | 13.65 | 1.30 | 0.03 | 0.55 | 0.91 | 0.02 | 0.01 | 0.03 |  |
| Brackish euc | 0.02 | 0.05 | 22.76 | 0.03 | 0.01 | 6.60 | 14.22 | 0.02 | 24.69 | 0.08 | 0.06 | 0.26 | 0.09 | x |
| Indoor peace lily | 0.05 | 0.08 | 2.87 | 0.04 | 0.01 | 2.39 | 0.90 | 0.04 | 0.77 | 0.24 | 0.00 | 0.01 | 0.09 |  |
| Town euc rain | 0.12 | 0.06 | 1.58 | 0.08 | 0.08 | 10.38 | 1.98 | 0.08 | 0.80 | 0.24 | 0.36 | 0.00 | 0.14 |  |
| Town euc no rain | 0.06 | 0.02 | 1.06 | 0.00 | 0.00 | 5.30 | 0.88 | 0.05 | 0.86 | 0.58 | 0.14 | 0.00 | 0.08 |  |
| Lake euc | 0.09 | 0.04 | 1.35 | 0.07 | 0.04 | 5.37 | 0.91 | 0.04 | 1.32 | 0.43 | 0.19 | 0.01 | 0.07 |  |
| Euc farm | 0.13 | 0.05 | 1.99 | 0.04 | 0.04 | 54.90 | 1.12 | 0.33 | 2.25 | 0.25 | 0.17 | 0.01 | 0.22 |  |

The water vapor sorption experiment results with the leaf wash samples are shown with standard error bars in Fig. 2. The percentage weight increase of moisture sorbed above the dry weight, Δ*w*, is plotted with relative humidity, RH, as calculated in Equation (3). Significant sorption occurs at high humidities at and above 75%RH with the ‘Brackish mangrove’ and ‘Cabinet mangrove’. Less so but still significant sorption occurs with the ‘Glasshouse chilli’ and the ‘Brackish euc’. All leaf washes appear to be somewhat hygroscopic. As the leaf sample washes are mixtures of different salts, minerals, oils, waxes and other materials, sorption behaves less predictably, especially above 84%RH and this is discussed further around Fig. 5. The two mangrove samples on the other hand behave more predictably due to their high NaCl content and minimal oils. As our focus is total water on the leaves, we consider the same data, plotted as the total of the triplicates, as total weight (mg) with relative humidity in Fig. A7, calculated with Equation (4). We compare the strongly deliquescent leaf wash samples that produced a clearly visible aqueous solution, in Fig. 3. The ‘Brackish mangrove’ as expected, was the most hygroscopic and becomes visibly liquid at 75%RH (POD of NaCl), aided by its high initial dry weight. The ‘Cabinet mangrove’ also sorbed large amounts of water and became visibly wet at 75%RH, compatible with the dominance of NaCl in the coating. The ‘Brackish euc’ became visibly wet at 84%RH, and the ‘Glasshouse chilli’ and ‘Ocean common reed’ at 97%RH. In Table 4 we compare these results to the ICP-OES data in Table 3 for the combined weight of Ca, Mg and Na in *μ*g (that are postulated to be the most hygroscopic). We see the five samples that visibly deliquesce in Fig. 3 correspond well to the five highest combined masses of these cations. The highest to lowest of the sum of Ca, Mg and Na also correspond reasonably well to the ranking of the maximum percentage weight increase over the dry weight, Δ*w* - %, in Fig. 2, which suggests that hygroscopicity of the leaf wash material may be predicted from the combined mass of Ca, Mg and Na from the ICP-OES data. For the ‘Brackish mangrove’, the materials present significantly deliquesced, over the total sample (total of three repeats), at 97%RH, produced 316 mg, as shown in Figs. 3. This equates to 0.03–0.05 mL on one leaf (assuming uniformity over 6–10 leaves), with a thickness of liquid on one leaf of 7.3 *μ*m (or 14.6 *μ*m considering salt glands are mainly on one abaxial surface), which is similar but higher than estimates of 1 *μ*m using other methods with plants that do not excrete salt [11]. We note other samples were hygroscopic and could also be deliquescent but were not included in Fig. 3 as they did not visibly deliquesce. Table 5 compares the dry weight and maximum sorption to the leaf area, ordered in the same way as Table 4. The two mangrove samples produce a similar dry weight and maximum sorption.

**Figure 2:**
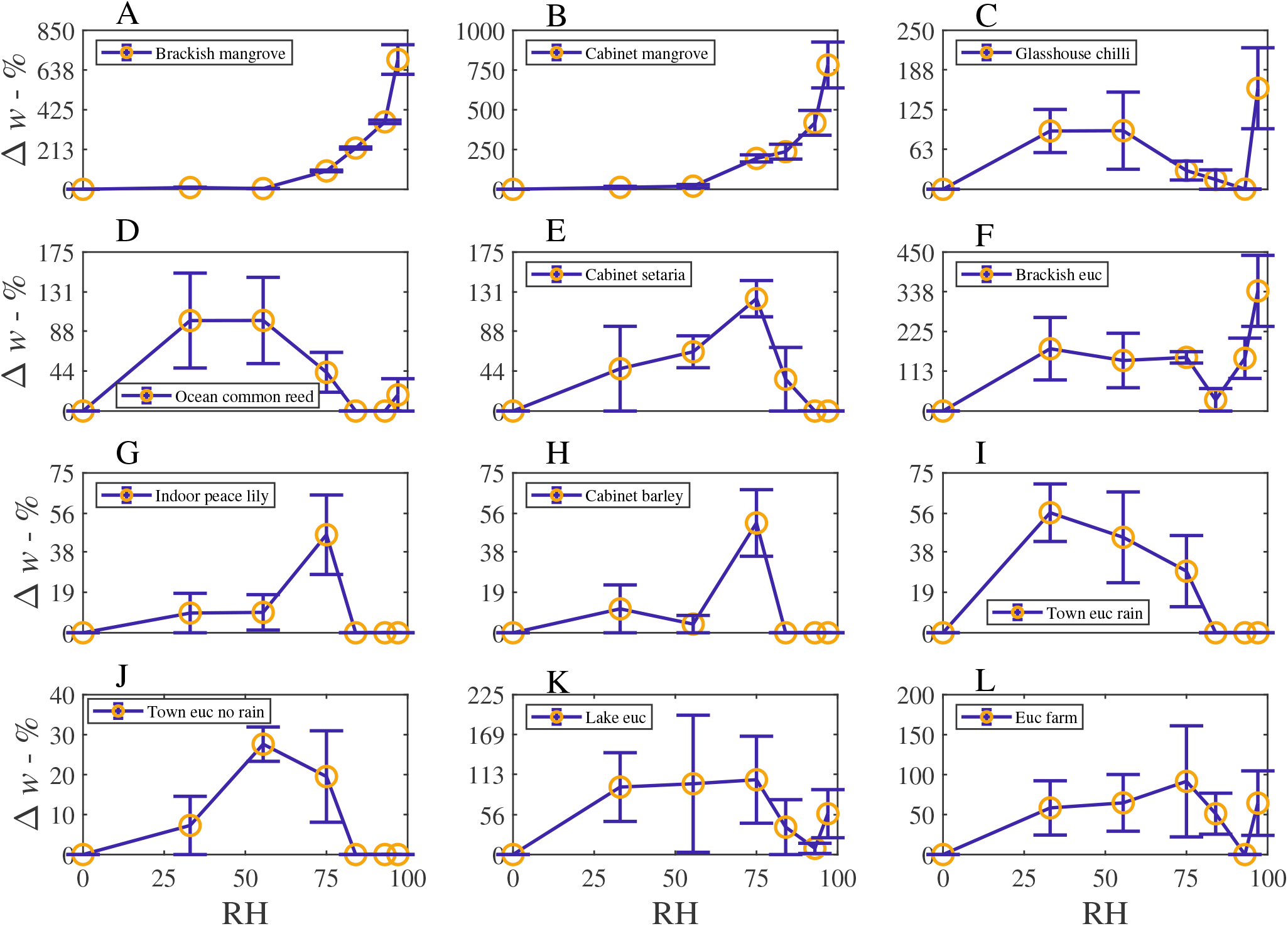
The percentage weight increase of moisture adsorbed of the leaf wash samples, Δ*w* - %, vs relative humidity, RH - %, plotted with standard error bars. Note the *x*-axis range is always the same but the *y*-axis range changes for each subfigure.

**Figure 3:**
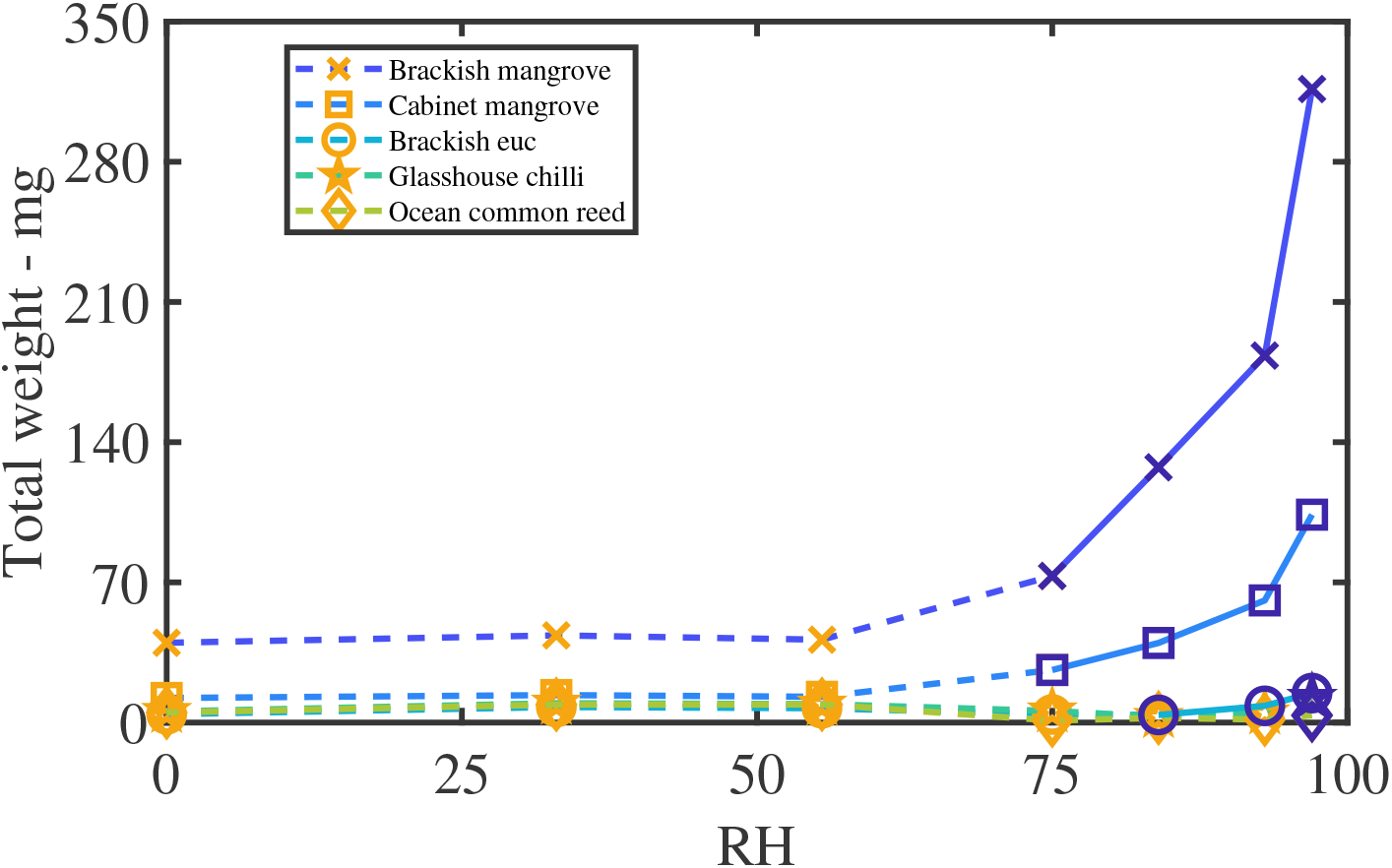
Samples that deliquesce visibly to the naked eye. Total weight (of three repeats with the dry weight, scaled with the blank) vs humidity (%). The orange symbols and dashed lines indicate water is not yet visible (but may have hygroscopic growth), and the blue symbols and solid lines indicate where an aqueous solution was clearly visible. Note the significant weight of the brackish mangrove sample at 97%RH of 316 mg.

**Table 4:**
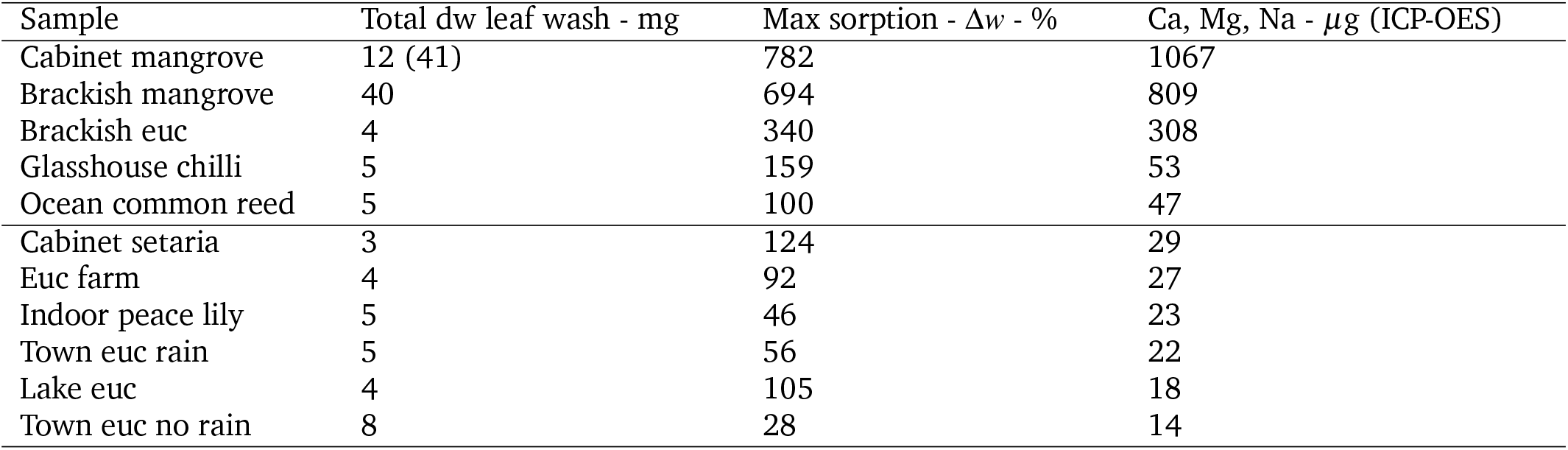
Total dry weight of each leaf wash sample (sum of the three repeats) after oven drying and centrifuging. The maximum sorption weight percentage increase over the dry weight Δ*w*, is compared with the combined weight in *μ*g of the Ca, Mg and Na ICP-OES data and the list is ordered by this combined weight. Samples above the center line visibly deliquesced. The combined *μ*g of Ca, Mg and Na has a positive correlation with visible deliquescence and maximum sorption. The dry weight may include some materials other than salts (such as oils, fine plant matter, waxes etc) present after centrifuging and increase the dry weight but not appear in the ICP analyses (for example’Town euc no rain’). Note ‘Cabinet mangrove’ comprised of only 3 leaves so the dry weight is scaled accordingly (as shown in brackets).

**Table 5:** Dry weight per leaf area and maximum wet weight per leaf area, in *μ*g cm^*−*2^. The cabinet mangrove sample comprised only 3 leaves so the scaled value from Table 4 has been utilized. The salt glands of the two mangrove species are located on the abaxial surface and as such are scaled by 2 (as shown in brackets). Samples above the central line are visibly deliquescent.

| Sample | dw per leaf area<br>$\mu\text{g cm}^{-2}$ | Max sorption per leaf area<br>$\mu\text{g cm}^{-2}$ |
| --- | --- | --- |
| Cabinet mangrove | 95 (190) | 805 (1609) |
| Brackish mangrove | 93 (185) | 736 (1471) |
| Brackish euc | 10 | 34 |
| Glasshouse chilli | 13 | 29 |
| Ocean common reed | 12 | 21 |
| Cabinet setaria | 6 | 10 |
| Euc farm | 9 | 23 |
| Indoor peace lily | 12 | 12 |
| Town euc rain | 12 | 19 |
| Lake euc | 9 | 14 |
| Town euc no rain | 18 | 23 |

Control experiments for salt and oil sorption were conducted, for comparison to the leaf washes, to better understand whether the results were similar when salt mixtures and oils were present, as shown in Figs. A8–A11. All the controls deliquesced and formed an aqueous solution that was visible to the naked eye, except the two pure oils - *Eucalyptus* and tea tree. The salts and mixtures with CaCl_2_ generally formed liquid at 32%RH (POD of CaCl_2_). Similarly mixtures with NaCl deliquesce at 75% (POD of NaCl). The mangrove nutrient (that the ‘Cabinet mangrove’ grows in a solution of) is dominantly made up of NaCl but also contains other salts with a lower POD. When comparing the mixture of NaCl plus *Eucalyptus* oil sample to the large NaCl sample, the POD is similar, however the change in weight is less when oil is included, indicating the oil was able to prevent some level of sorption but did not prevent deliquescence. When applying this outcome to the leaf wash results, it could be relevant for the ‘Brackish euc’, which was able to form an aqueous solution, despite the presence of some oils but to a lesser extent than the mangroves (Fig. 3). We note that the small NaCl sample has higher than expected sorption at 33%RH and this might be corrected with additional repeats.

The experimental data in Fig. 4, for the ‘Brackish’ and ‘Cabinet mangrove’, compared to the mangrove nutrient and NaCl control, is modeled with the Guggenheim, Anderson, and de Böer (GAB) isotherm [3–5] (Equation (A5)). Fig. 4 shows that the percentage moisture gain is similar for both mangroves, the GAB isotherm has a similar trend, and the percentage sorption is very similar at nearly all humidities. The sorption of the two mangrove samples compare well to the NaCl control and mangrove nutrient (mostly NaCl), and therefore the sorption is likely driven largely by NaCl. The ‘Brackish’ and ‘Cabinet mangrove’ sorption at humidities between 75–100%RH, is somewhat higher than the mangrove nutrient and NaCl, which indicates that additional hygroscopic compounds are present, for example CaCl_2_. When comparing the percentage moisture gain to the total weight in Fig. 3, the ‘Brackish mangrove’ has a much greater total weight at 97%RH but also started out with a heavier dry weight. When scaled with the dry weight, the percentage moisture gains of both mangrove samples are similar (Fig. 4). The GAB constants have physical meaning and the values of *k* for the four isotherms are similar, so the interaction energy between the multiple layers of water is similar. The values for *w*_*s*_ are higher for the two controls, indicating that they have a larger monolayer saturation value but possibly less multilayers than the two leaf washes. In summary, the two mangroves studied from a cabinet and near brackish water, both excrete salt through the leaves, have sorption properties similar to, but greater than, NaCl and the mangrove nutrient, suggesting a dominance of NaCl but also with other hygroscopic materials.

**Figure 4:**
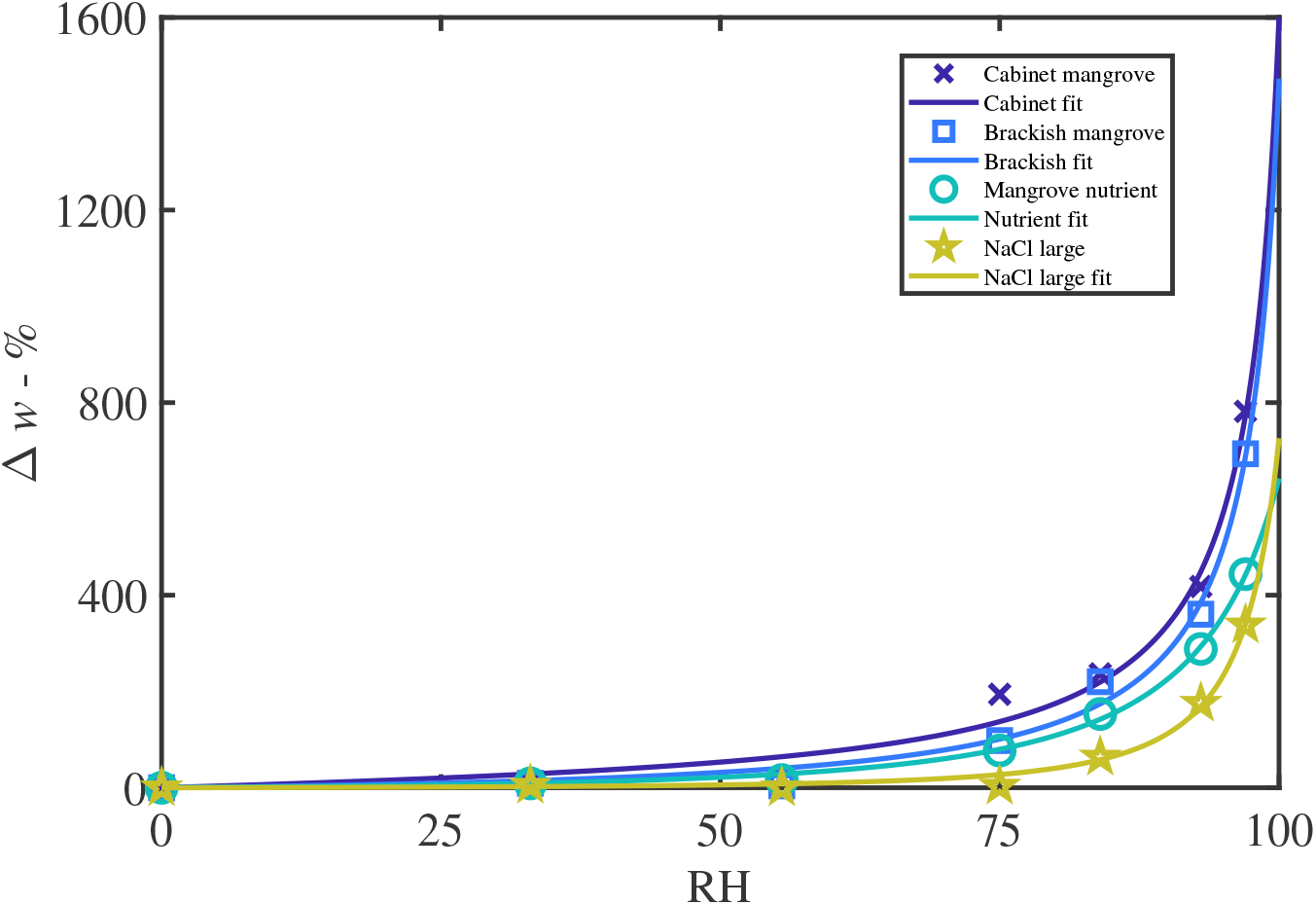
A selection of adsorption isotherms including cabinet and brackish mangroves, along with the controls of the mangrove nutrient and NaCl with a large sample size. The plot is percentage moisture gain over the dry weight vs relative humidity. The data are fitted with the GAB isotherm, as described in Equation (A5). The controls were formulated with a similar dry weight to the leaf wash samples. The parameters are described in Table A7 and R^2^ values are greater than 98.6%.

We found a significant presence of lipophilic compounds, possibly oils or waxes. The presence of *Eucalyptus* oils in some leaf wash samples is indicated by the silver nitrate study, shown in Fig. A6, and waxes were found with the XRD (for example Fig. A15). In Fig. 5, as leaf washes are mixtures of salts and lipophilic compounds, the sorption of the samples is less predictable. In Fig. 5, by comparing the *Eucalyptus* leaf wash samples to the *Eucalyptus* and tea tree oil controls, the presence of oils in the leaf wash samples results in increased sorption followed by a decrease at high humidity, as the oils dominate the signature, similar to the pure oils. The ‘Brackish euc’ is able to overcome the effect of the oil at high humidity and increased sorption is possible, to the point of visibly deliquescing. ‘Brackish euc’ has the large presence of salts comparing *Eucalyptus* samples (*μ*g of Ca, Mg and Na in Table 4). The sorption overcomes that of the oil after a critical mass of water is reached, the salts dissociate, and high sorption occurs. The ‘NaCl with euc oil’ control sample in Fig. A8 (J), indicates a similar behavior.

**Figure 5:**
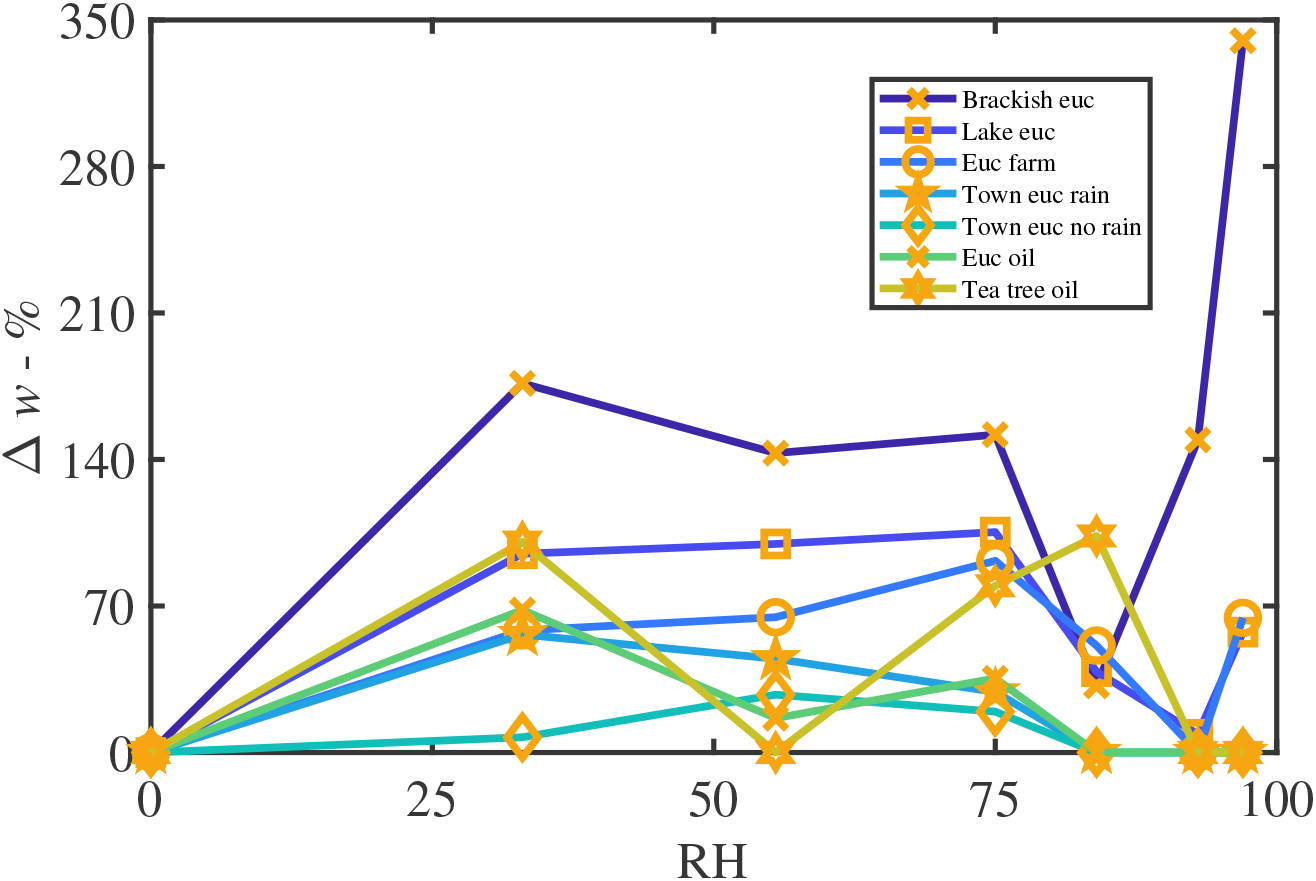
Percentage weight gain of moisture vs relative humidity of all *Eucalyptus* leaf wash samples, along with two oil controls for comparison. The oils contributed to the increasing sorption at low humidity then decrease in sorption between 33%RH and 84%RH, while the other hygroscopic ionic compounds present in the sample contributed to weight gain at high humidities such as shown by ‘Brackish euc’. The oil controls follow a similar trend to the leaf washes.

## 4. Discussion

Materials including atmospheric aerosols on leaf surfaces substantially impact on the interaction between the leaf surface and atmospheric moisture, even for plants grown in comparatively clean environments (for example growth cabinets and glasshouses). Our results, in line with other recent work [27, 28], show that not only microscopic amounts of water but relatively large volumes of water visible to the naked eye can form on plant leaf surfaces. This water can influence stomatal function and anatomy [27, 28]. Our results suggest that some experimental designs may require the incorporation of tests, to determine if leaf washing and air filtration are necessary and if a range of relative humidities is required instead of a single humidity, depending on the research question. These considerations may be particularly important when comparing crops/field work to glasshouse/cabinet studies, agrochemical penetration experiments with foliar sprays, foliar water uptake studies and certain gas exchange experiments. We note the surprising case of the sorption of ‘Glasshouse chilli’ that was able to form water visible at 97%RH. This plant was two years old, did not experience any leaf washing. It is demonstrably possible that plants grown in a range of locations from glasshouses to growth cabinets may experience this phenomenon, if washing of their leaves does not occur over prolonged periods. The ‘Town euc rain’ sample contains impurities present in rain with larger amounts of Al, Cu, Fe, Si and Zn present in the ICP-OES data in Table 3. ‘Brackish euc’ has a larger than expected amount of Ca, unexpected as it is higher than the adjacent ‘Brackish mangrove’ plant. The majority of the dry weight of both mangroves originates from the interior of the plant excreted through salt glands, though some aerosols and saltwater spray exist on the ‘Brackish mangrove’ sample. The ‘Euc farm’ sample has greater levels of K, Al, Mn and Zn detected in the ICP-OES data, likely from agrochemical spray drift or particles from the nearby highway. Some samples in Table 3 including ‘Euc farm’ had large amounts of K present but were usually associated with *Eucalyptus* hence had reduced sorption caused by the presence of oils.

Hygroscopic materials on the plant leaf surface will be affected by the adjacent/local humidity. Due to the boundary layer [11], temperature and the action of stomata releasing water vapor from the interior of the leaf, the humidity on the leaf surface may be higher than the ambient relative humidity in the air. Comparing air to leaf surface humidity, an increase of 35%RH above the daytime ambient humidity has been found [4]. Applying this 35%RH increase, for example if the ambient humidity is 40% and the leaf surface humidity is at 75%, then NaCl (POD 75%RH) on the leaf surface will be able to deliquesce and form liquid water. When suspended or on a passive surface, such as rock, the humidity would be below the POD of NaCl and closer to the ambient humidity (of 40%), thus the NaCl cannot deliquesce. When applying this boundary layer effect to sorption of salts *in situ* on the plant leaf, salts will sorb moisture from the local environment even though the ambient relative humidity is less than the salts POD. Due to the boundary layer, salts on plant leaves may form an aqueous solution at a much larger range of ambient humidities, further increasing the significance of this study. The boundary layer is relevant to plant leaves in their natural environment, but may have less importance in artificial experimental settings, for example the cuvette of a photosynthesis system (for example Li-6400 or Li-6800) where a high fan speed is used to minimize the boundary layer thickness and in an environmental scanning electron microscope (ESEM) the vacuum conditions would minimize the boundary layer, therefore results of salts on leaves viewed with ESEM may not be influenced by this boundary layer effect.

From observations during the salt control experiments, 1 g of solid CaCl_2_ can sorb significant proportions of water and visibly deliquesce very quickly, in a matter of 5–10 mins in relatively low humidities, while it may take days to reach adsorption equilibrium, or completely dry out again in the oven (desorption). We note that in the context of hygroscopic materials on plant leaves in the environment, these adsorption and desorption timescales are very relevant as temperature and humidity change throughout the day. For example, if the air humidity increases, water may be quickly adsorbed by these materials, or if the air temperature increases in the morning leading to a decrease in air humidity, hygroscopic particles on the leaf surface will extend evaporation times of residual water, and keep the leaf wet longer. These adsorption and desorption timescales for hygroscopic particles *in situ* on the plant leaf require further research. The experimental data of the current work of sorption of materials on leaves, Δ*w*, can be utilized in a mechanistic model [47] for droplet evaporation. If considering evaporation where the RH is changing significantly with time, over the evaporation timescale, it may be necessary to consider additional mechanisms [29] including desorption hysteresis (involving crystallization or the point of efflorescence, for example NaCl [43]) of the material on the plant leaf.

Future work could include desorption, sorption data measured with time increments over a time span of longer than 3 days, conducting a similar study but looking more specifically at one location including many plant species, and individuals/biological replicates within that species. Water vapor sorption experiments could be performed on the leaf wash pellet left behind after centrifugation, which may contain insoluble waxes and mineral grains, organic debris, insect material and material sloughed from the leaf itself. This is relevant for the brackish mangrove sample, where the leaves had a larger amount of dried mineral dust present as they were periodically submerged at certain tides. The pellets were tested via XRD for the same species as in Figs. A12–A13 but no NaCl or other salts were found, and were not investigated further.

Six-week-old spongy tobacco leaves (native Australian tobacco (*Nicotiana benthamiana*)) were deemed unsuitable for experimentation as when shaken the leaves stained the water green, likely with chlorophyll leaching from ruptured cells. We note that it is uncommon for sorption isotherms of oils to be studied. When studied, stabilizers are often included and sorption experiments on oils can take 1–2 weeks to reach equilibrium [15]. Further research is required, especially for prolific *Eucalyptus*. Hygroscopic salts on leaves assist in maintaining leaf surface wetness when stomata are open during the day even at low humidity, maintain stomatal function and development, assist with foliar water uptake and extend the droplet evaporation time of dewfall. In terms of foliar applied agrochemicals, hygroscopic salts on the leaf either *in situ* or in the applied droplet formulation, can impact agrochemical penetration by altering the formulation’s effectiveness at a given relative humidity, rate of droplet evaporation, penetration effectiveness of active ingredients and surface tension. Salts can also impact the experimental setup and design, and if leaves are regularly washed with water or not, especially when comparing field and glasshouse grown plants. As the surface properties of leaves vary a great deal, as do environmental properties, no single study can be specifically applicable to other plants and areas. Despite this, the general cases explored here have wide-reaching implications in ecological, physiological and agricultural studies.

This work has demonstrated our aim of showing that the materials on plant leaves are hygroscopic and that an aqueous solution can form over small amounts of particles on the plant leaf surface grown in a range of environments. Five leaf washes attracted water to the point of visibly deliquescing, even in a glasshouse plant. Mangroves that excrete salt are covered with a layer that can form up to a total of 0.3 mL of liquid (for 6–10 leaves or 30 *μ*L on one leaf) at high humidities. This salt is mostly NaCl, but also contains other hygroscopic particles. An unexpected outcome of our study was the higher than expected levels of oils on the surface for all *Eucalyptus* samples studied.

## Funding

The authors acknowledge funding provided by the Australian Research Council Centre of Excellence for Translational Photosynthesis (CE1401000015).

## Acknowledgment

The authors thank Ulrike Troitzsch (ANU) for conducting the XRD experiments, Brett Knowles (ANU) for assisting with the ICP-OES experiments and Graham Farquhar (ANU) for his advice and thoughtful conversations.

## Author contributions

ET is responsible for project motivation, experimental design, conducting vapor sorption experiments, analysis, model fitting, results and article writing, editing and revision. HS is responsible for the experimental and biological consultation, article editing and revision. TE is responsible for conducting ICP-OES experiments and methodology, article editing and revision.

## Conflict of interest statement

The authors declare that the research was conducted in the absence of any commercial or financial relationships that could be construed as a potential conflict of interest.

## Appendix

**Table A6:** Salt and oil control mixtures. Samples were dissolved/suspended in de-ionised water and oven dried. The percentage mix is found by weighing the salts before creating the solution. For all the oils, approximately 100 mg was the wet weight before oven drying, most of the oil evaporated in the oven. The mangrove nutrient is mainly NaCl, along with 1295 ppm Mg, 430ppm Ca and 390 ppm K.

| Compound | Total dry weight - mg | Percentage mix - % |
| --- | --- | --- |
| Mangrove nutrient | 21.6 | - |
| NaCl + CaCl<sub>2</sub> | 60.0 | 41 NaCl, 59 CaCl<sub>2</sub> |
| NaCl + CaCl<sub>2</sub> + KCl | 172.5 | 12 NaCl, 71 CaCl<sub>2</sub>, 17 KCl |
| CaCl<sub>2</sub> + KCl + NaCl + Ca(NO<sub>3</sub>)<sub>2</sub> | 96.5 | 37 CaCl<sub>2</sub>, 11 KCl, 17 NaCl, 35 Ca(NO<sub>3</sub>)<sub>2</sub> |
| Ca(NO<sub>3</sub>)<sub>2</sub> + KCl | 105.3 | 11 Ca(NO<sub>3</sub>)<sub>2</sub>, 89 KCl |
| NaCl + euc oil | 49.0 | 96 NaCl, 4 euc oil |
| NaCl large | 24.4 | - |
| NaCl small | 1.1 | - |
| CaCl<sub>2</sub> large | 35.8 | - |
| CaCl<sub>2</sub> small | 2.1 | - |
| Euc oil | 2.7 | - |
| Tea tree oil | 0.9 | - |

### A1. Chloride and Sulphate Presence Test

To test for the presence of chloride (Cl^−^) and sulphate (SO_4_^2 –^), the most abundant anions in sea-water[1], 2 mL of liquid was removed from the 60 mL leaf sample washes destined for the sorption experiment. 1 mL aliquot of solution placed in clean 2 mL glass vials was used to identify chloride and sulphate non-quantitatively. For chloride, a few drops of dilute nitric acid followed by a few drops of silver nitrate (AgNO_3_) in solution (2% (w/v)); were added to the 1 mL sample. After 10 minute the presence of a precipitate was recorded. The silver nitrate test had the unexpected effect of turning some samples yellow/brown, indicating the presence of oils such as *Eucalyptus* oil and silver nanoparticles may have caused this coloration [2]. Ag^+^ is known to react with lipopolysaccharides and amino acids. For the sulphate test, the same procedure was conducted but a few drops of barium chloride solution (2% (w/v)) followed after 10 minutes by a few drops of dilute hydrochloric acid.

### A2. GAB isotherm

We consider the Guggenheim, Anderson, and de Böer (GAB) isotherm[3–6] for fitting a selection of samples. The GAB isotherm describes water adsorption as a monolayer that can form multilayers at high humidities. It is formulated by applying the Langmuir isotherm to each layer (evaporation and condensation can occur only from or on exposed surfaces) and is at equilibrium where the rate of adsorption equals the rate of desorption. The GAB isotherm is as follows:

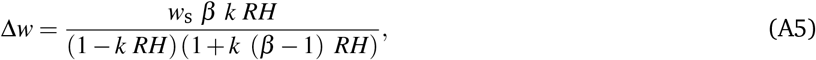

where Δ*w* is the percentage weight increase above dry weight at each relative humidity as a percentage, *RH* (*RH* = *a*_*w*_ × 100% = *p/p*_0_ × 100%), *w*_S_ is the monolayer of water adsorbed per solid, *β* is the equilibrium parameter of adsorbed water or the interaction energy between water and solid and *k* represents the difference in free enthalpy of the water molecules in the monolayer and layers above the monolayer, where *β* and *k* depend on temperature with the Arrhenius equation and *k* is always less than 1. The fitting and analysis were performed using MATLAB^®^ (Mathworks, U.S.A.).

**Table A7:**
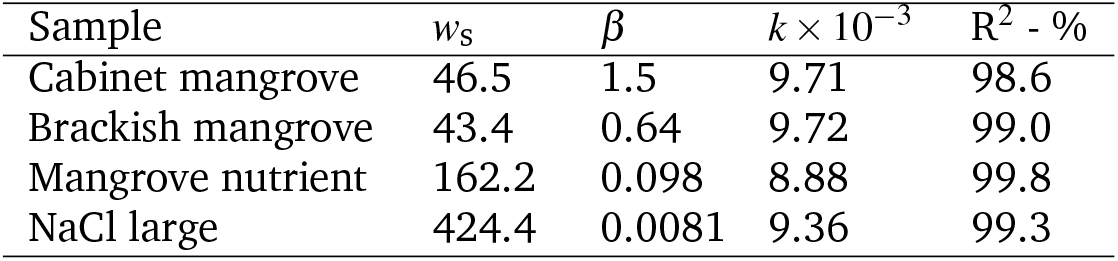
GAB parameters for Figure 4, fitted with Equation (A5)

| Sample | $w_s$ | $\beta$ | $k \times 10^{-3}$ | R<sup>2</sup> - % |
| --- | --- | --- | --- | --- |
| Cabinet mangrove | 46.5 | 1.5 | 9.71 | 98.6 |
| Brackish mangrove | 43.4 | 0.64 | 9.72 | 99.0 |
| Mangrove nutrient | 162.2 | 0.098 | 8.88 | 99.8 |
| NaCl large | 424.4 | 0.0081 | 9.36 | 99.3 |

**Table A8:** Operating parameters for ICP-OES measurements.

| Parameter | Value |
| --- | --- |
| Read time | 20 s |
| Replicates | 3 |
| Sample uptake delay | 25 s |
| Rinse time | $30 \times 2$ s (0.5 M HNO<sub>3</sub> & MQ) |
| Stabilization time | 10 s |
| Pump speed | 12 rpm |
| Fast pump | ON |
| RF power | 1.5 kW |
| Aux flow | 1.0 L min<sup>-1</sup> |
| Plasma flow | 12.0 L min<sup>-1</sup> |
| Nebulizer flow | 0.7 L min<sup>-1</sup> |
| Viewing mode | SVDV |
| Viewing height | 5 mm |
| Background correction | FACT |
| Number of pixels | 2 |
| Quality control | 1 µg g<sup>-1</sup> multi element |
| Analytes | Al (396.152 nm); B (249.772 nm); Ca (422.673 nm); Cu (327.395 nm); Fe (238.204 nm); K (766.491 nm); Mg (279.553 nm); Mn (257.610 nm); Na (589.592 nm); P (213.618 nm); Si (251.611 nm); Sr (407.771 nm); Zn (213.857 nm) |
| Drift correction | C (193.027 nm); Ar (565.070 nm, 645.918 nm, 675.283 nm, 699.217 nm, 704.096 nm) |

**Figure A6:**
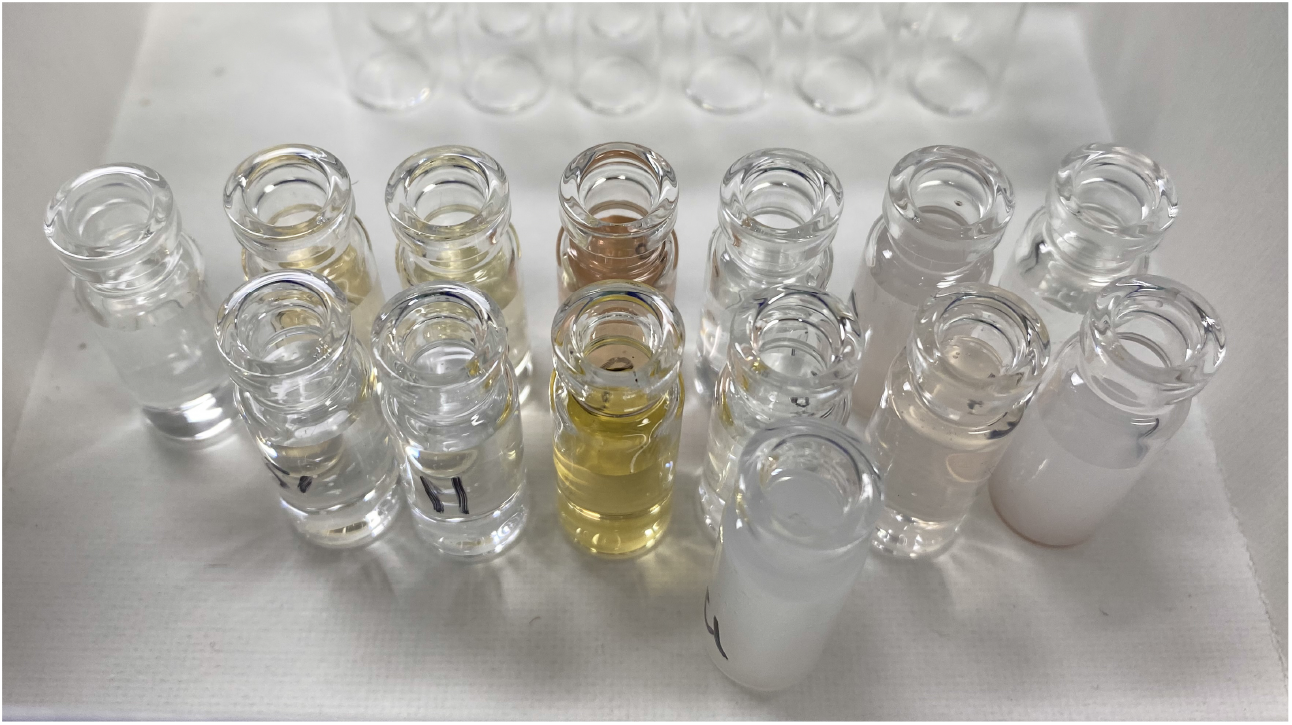
Silver nitrate test for the presence of Cl^−^. Samples that reacted formed a milky white precipitate of AgCl. These were Brackish mangrove, Brackish Eucalyptus and Cabinet mangrove. This test also showed the presence of nanoparticles related to Eucalyptus oils with a change in colour from clear of milky to yellow to brown. All 5 Eucalyptus samples changed colour. The darkest colour (copper brown) was Town euc no rain, the second was Town euc rain (dark yellow). The vial in the foreground is the CaCl_2_ control.

---

**Figure A7:**
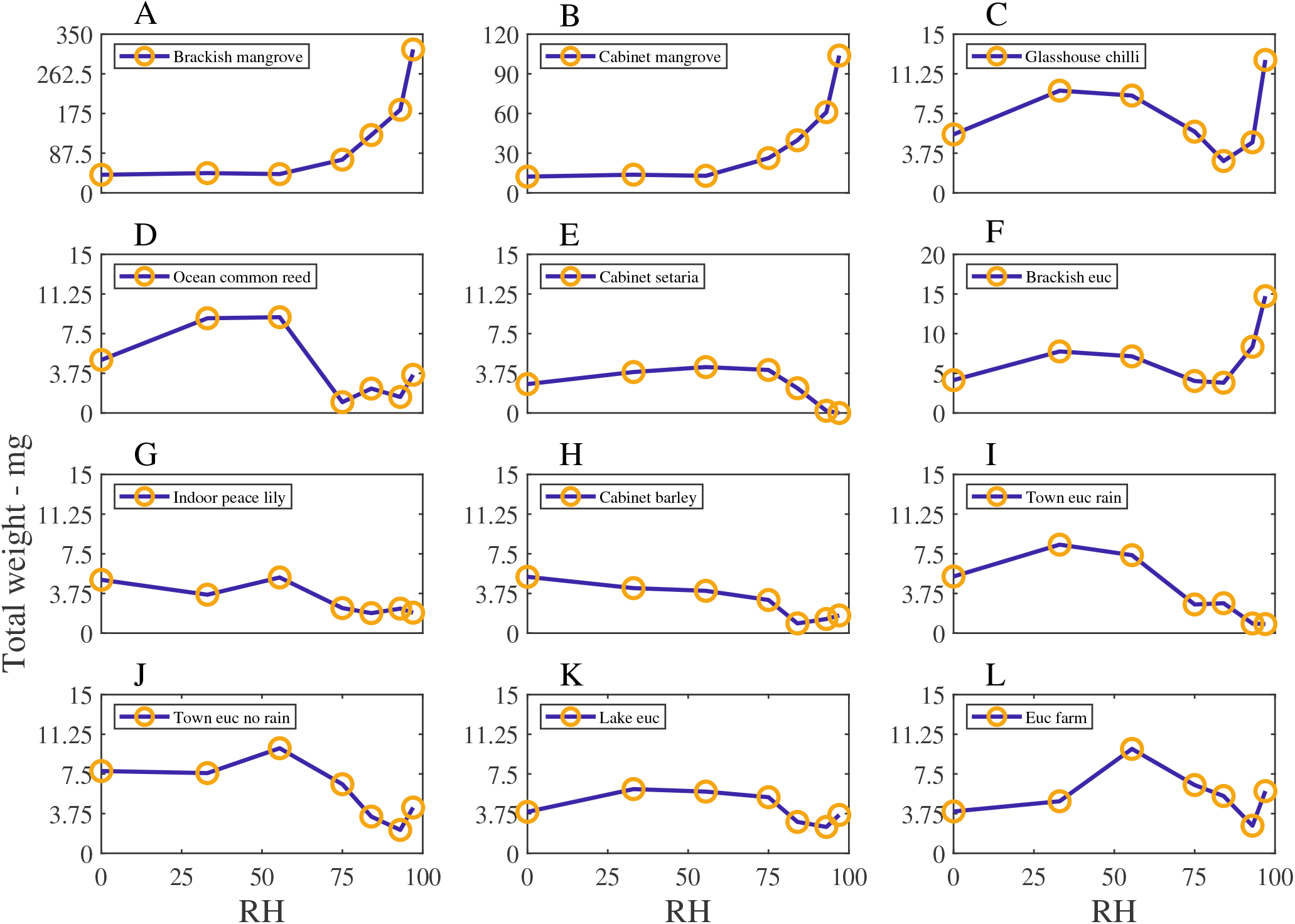
The Total weight (mg) of moisture adsorbed for the leaf wash samples, with relative humidity, RH (%). Total weight includes the dry weight and is scaled with the blank. Note that the *y*-axis range changes for each subfigure.

**Figure A8:**
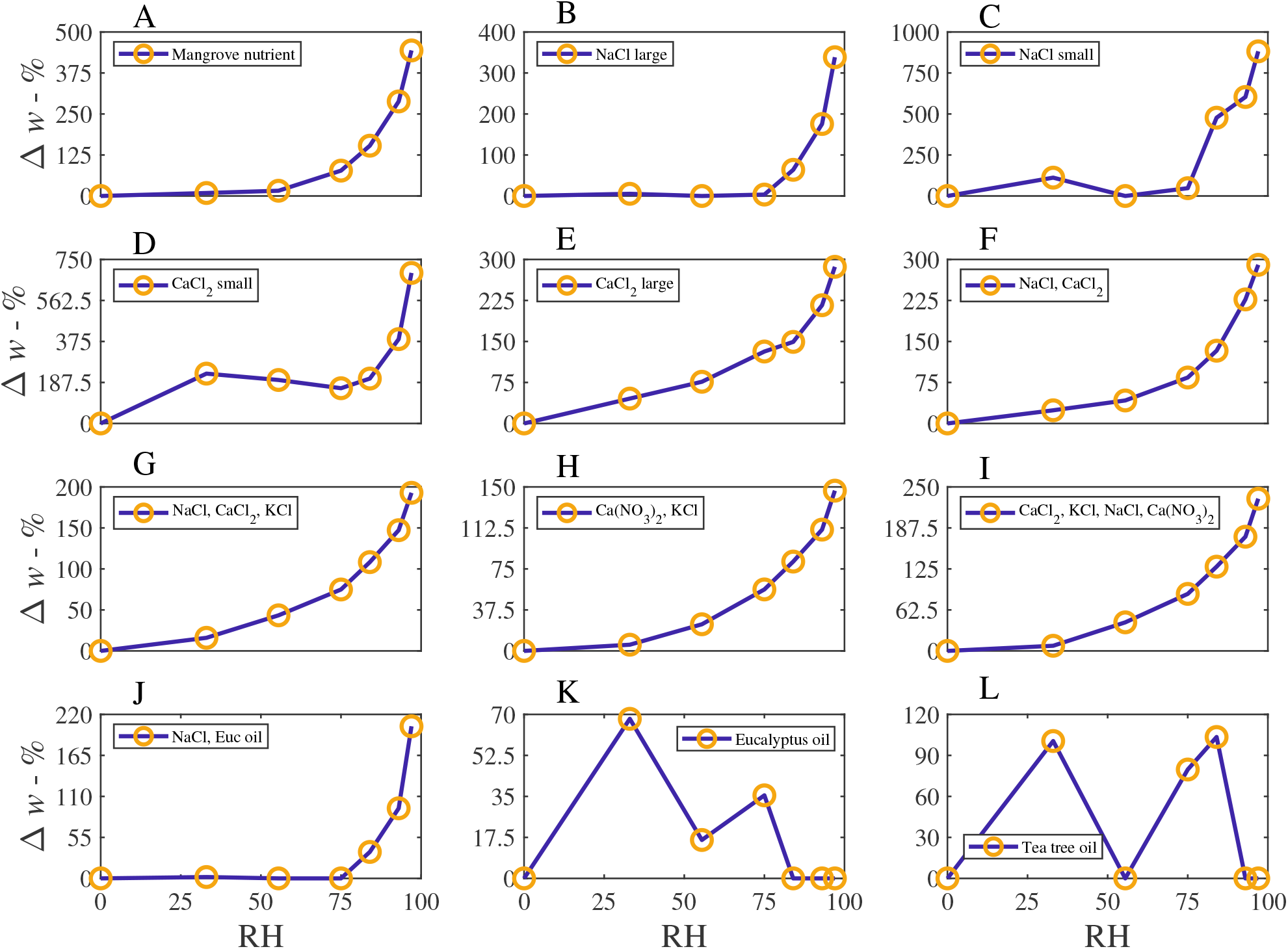
The percentage weight increase of moisture adsorbed above the dry weight of the controls, Δ*w* - %, with relative humidity, RH - %. The *y*-axis range changes for each subfigure.

**Figure A9:**
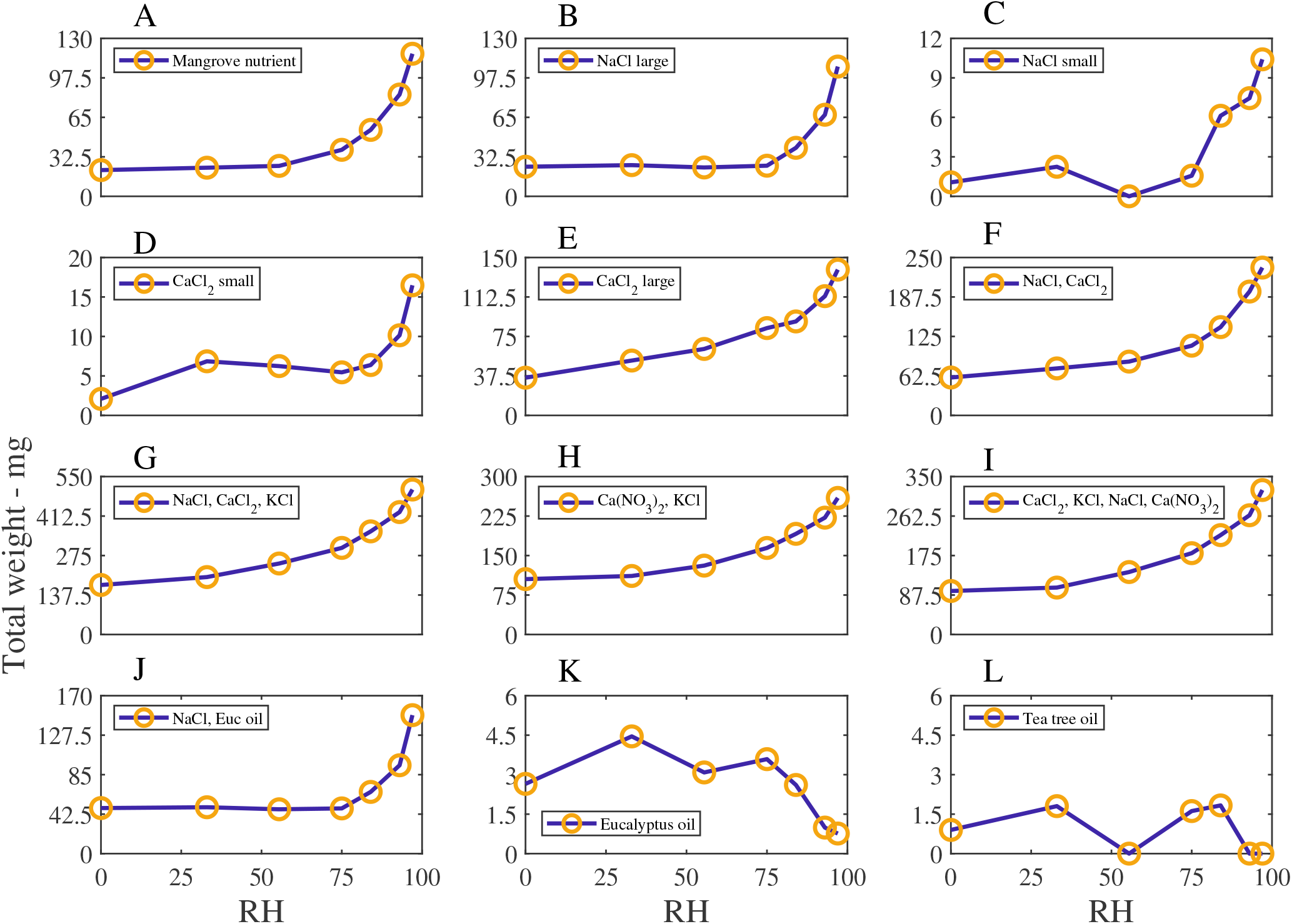
The Total weight (mg) of moisture adsorbed for the controls, with relative humidity, RH (%). The total weight includes the dry weight and is scaled with the blank. The *y*-axis range changes for each subfigure.

**Figure A10:**
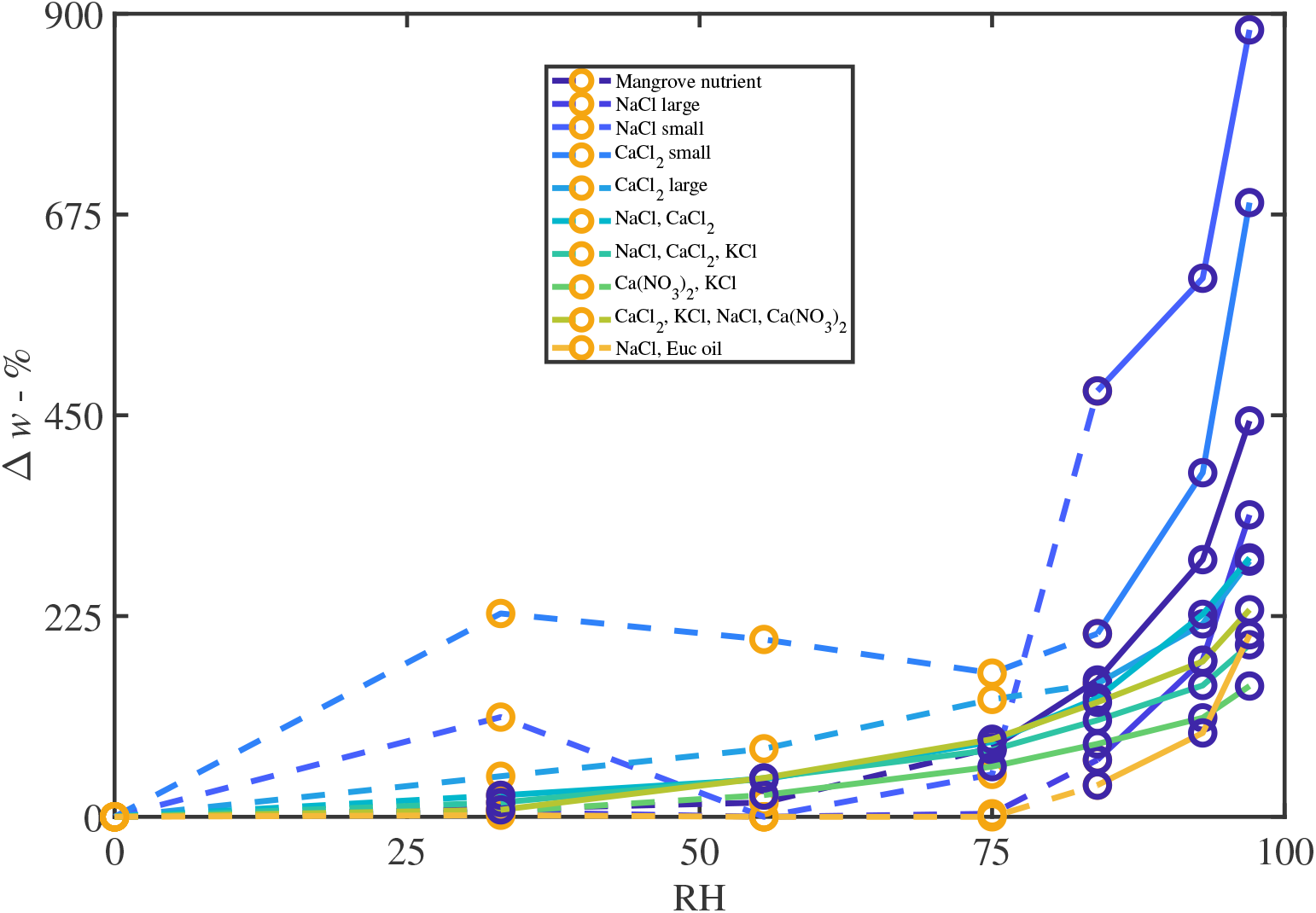
Control samples that have developed a coating of liquid water that is visible without the need for special equipment. The percentage weight increase of moisture adsorbed above the dry weight of the controls, Δ*w* - %, with relative humidity, RH - %, is shown. The orange circles and dashed lines indicate water is not yet visible, and the blue circles and solid lines indicates where liquid water is visible.

**Figure A11:**
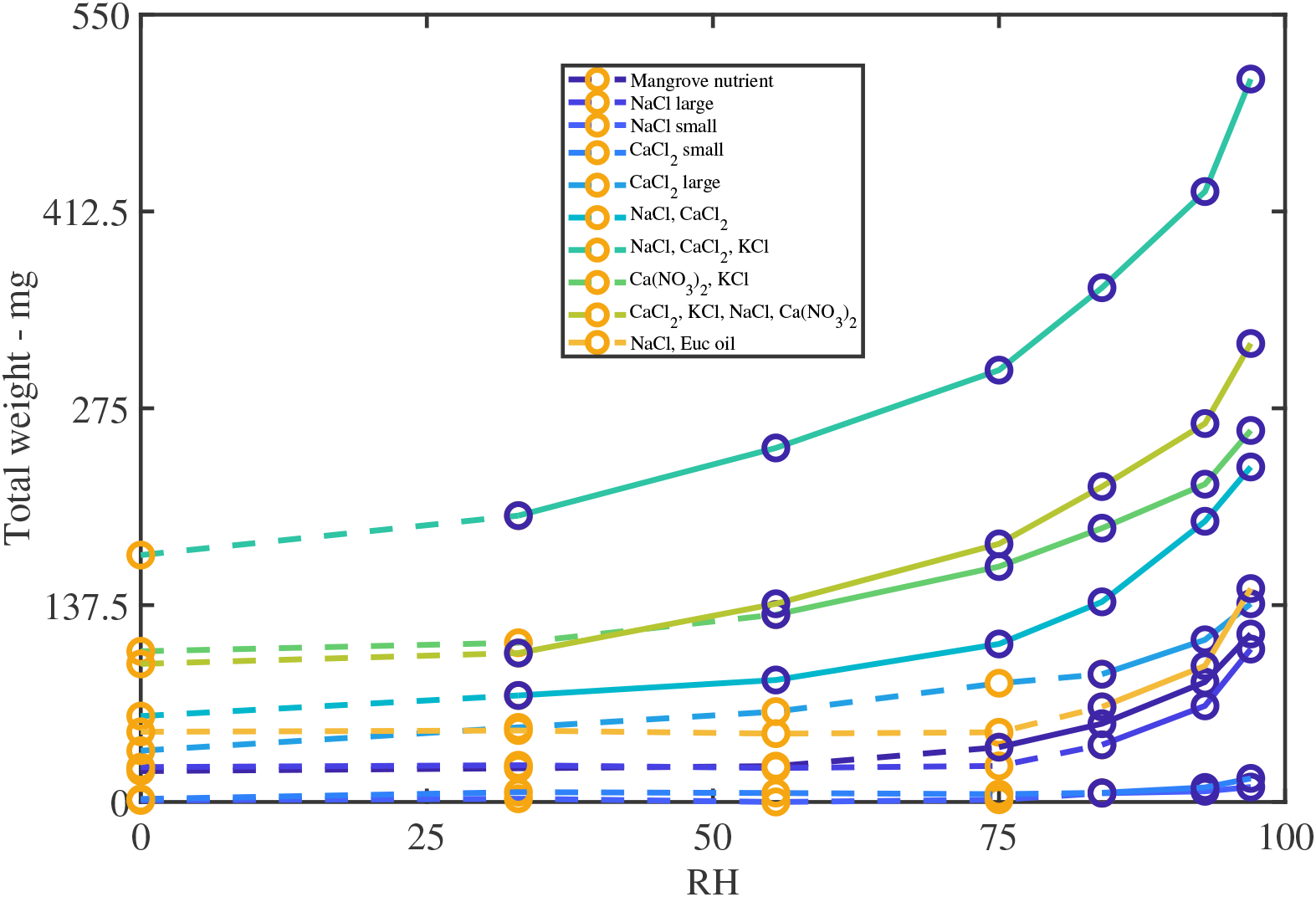
Control samples that have developed a coating of liquid water that is visible without the need for special equipment. The Total weight (mg) of moisture adsorbed for the controls, with relative humidity, RH (%), is shown. The orange circles and dashed lines indicate water is not yet visible, and the blue circles and solid lines indicates where liquid water is visible. Noteworthy is the significant weight of the NaCl + CaCl_2_ + KCl sample at 97%RH of 500 mg or 0.5 mL, although this sample started with the largest dry weight.

**Figure A12:**
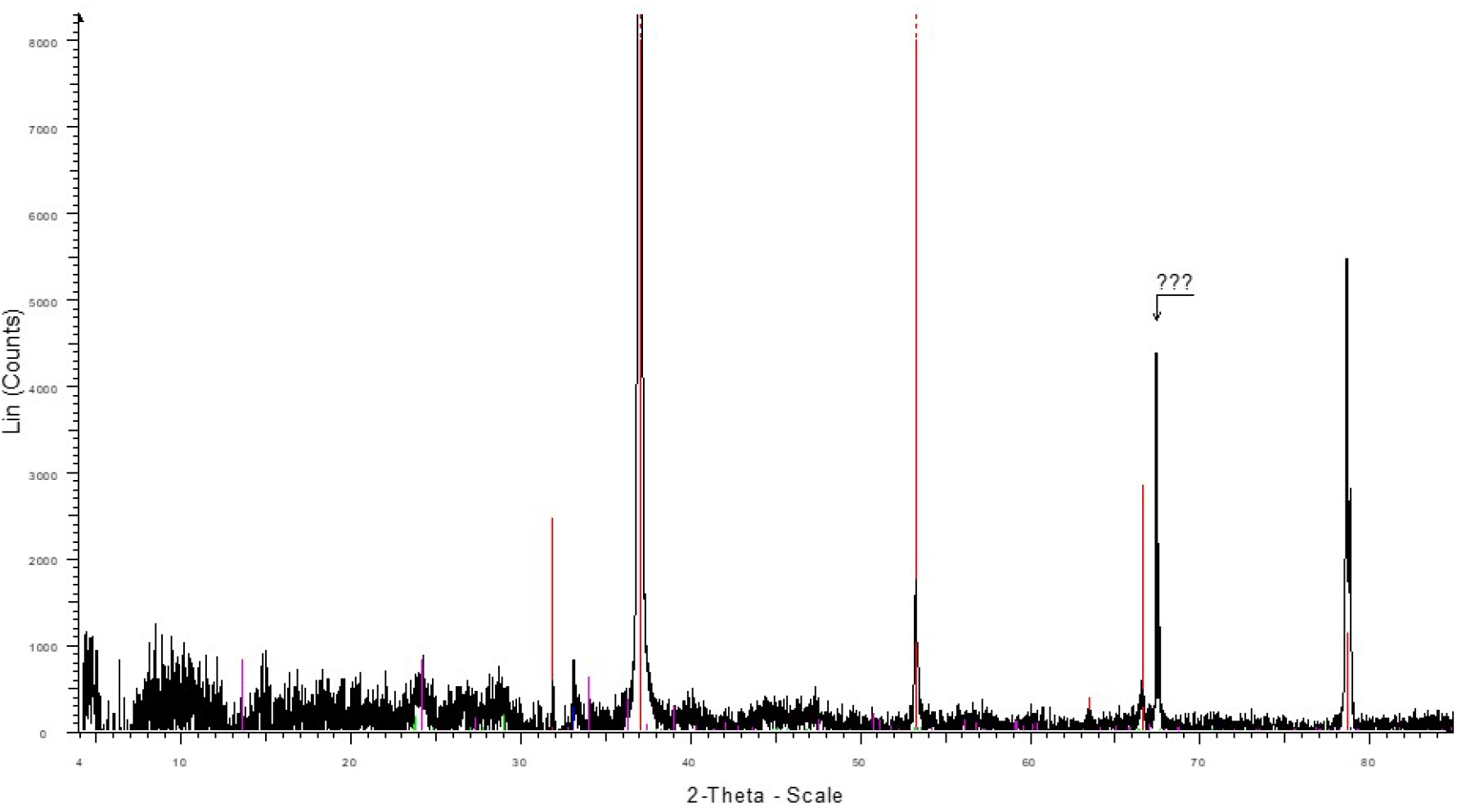
XRD results for Brackish mangrove leaf wash from the liquid wash, and corresponding results are shown in Table 2, with intensity (counts) vs angle (deg 2theta). Red is halite NaCl, blue is sylvite KCl, green is kaolinite Al_2_Si_2_O_5_(OH)_4_ and magenta is gypsum CaSO_4_. The question marks refer to an unidentified phase, possibly quartz.

**Figure A13:**
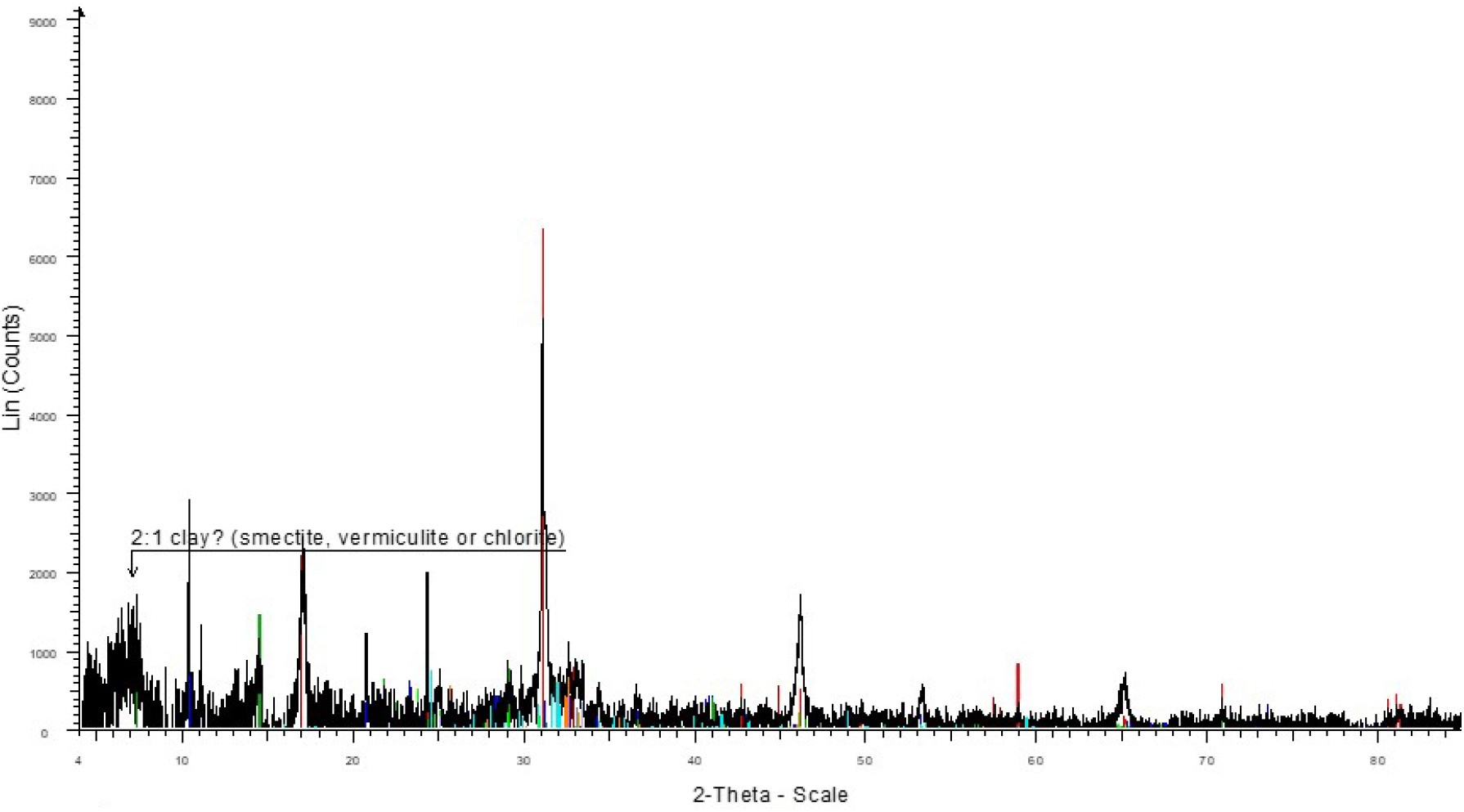
XRD results for Town euc rain leaf wash from the liquid wash, and corresponding results are shown in Table 2, with intensity (counts) vs angle (deg 2theta). Red is quartz SiO_2_, blue muscovite, light green is kaolinite Al_2_Si_2_O_5_(OH)_4_, dark green is chlorite, orange is plagioclase (Ca,Na)_1−2_(Si,Al)_2−3_O_8_, turquoise is K-feldspar KAlSi_3_O_8_, lilac is talc Mg_3_(OH)_2_Si_4_O_10_, grey is sylvite KCl and dark red is boehmite AlOOH.

**Figure A14:**
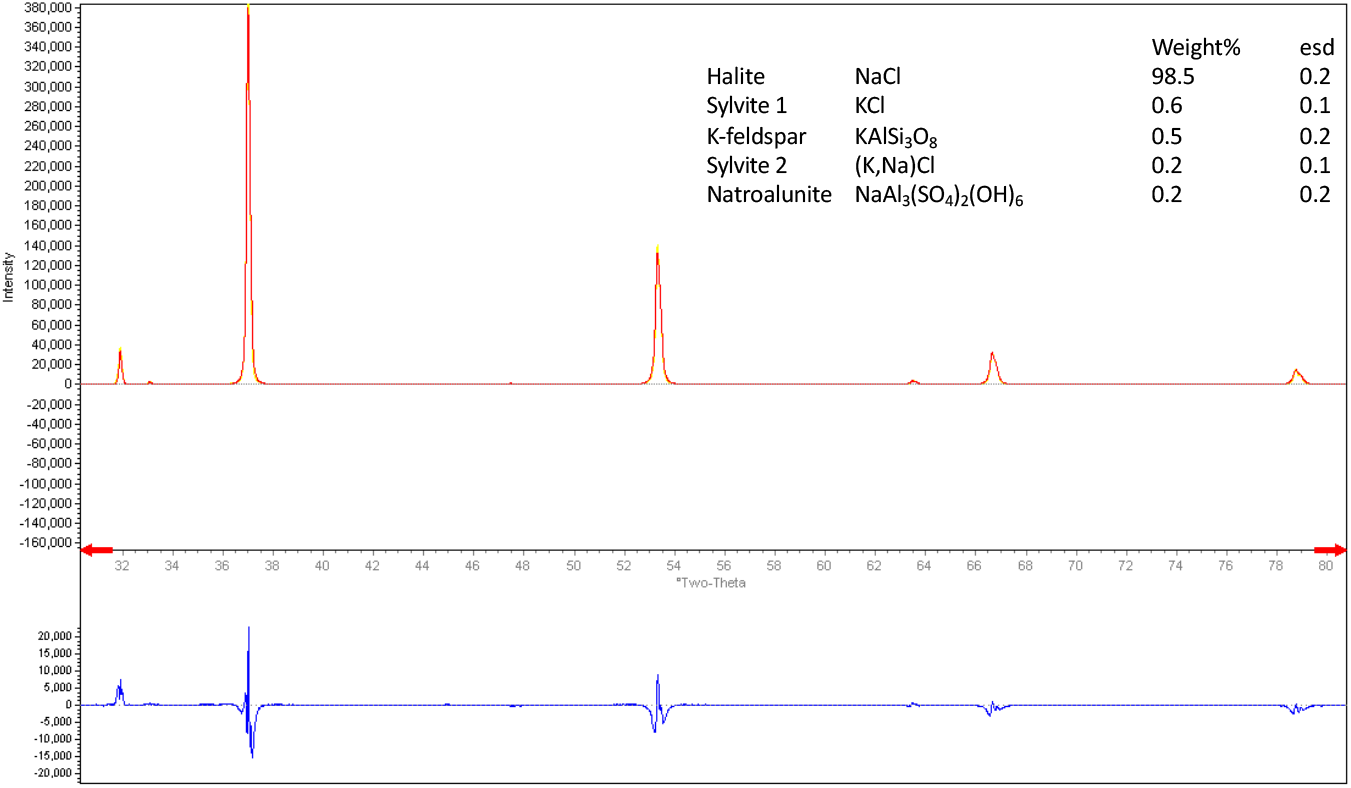
XRD results for a glasshouse mangrove, with intensity (counts) vs angle (deg 2theta). The sample for the XRD was collected by scraping the particles on the bottom of the leaf directly. Note this is the same plant species but a different individual as the Cabinet mangrove sample, and collected at a later date and grown in a glasshouse. No washing of the leaves occurred while the plant was growing.

**Figure A15:**
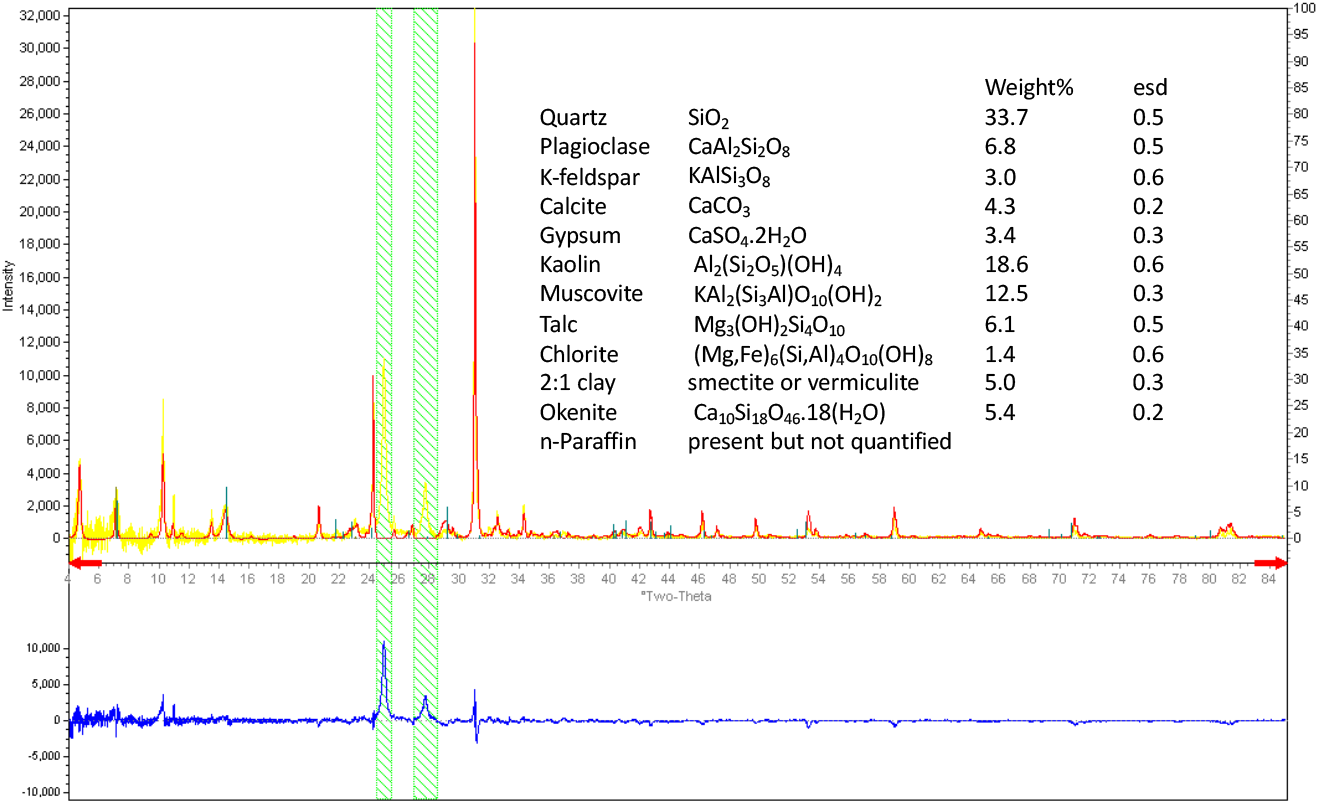
XRD results of Queensland kauri pine (*Agathis robusta* (syn. *A. palmerstonii))*, located indoors, with intensity (counts) vs angle (deg 2theta). This species was not included in the original sample set but was located in the same space as the Indoor peace lily. The plant was mature and had very waxy leaves that had a whitish sheen. Several waxes were seen but not quantified (n-Paraffin). The sample for the XRD was collected by scraping the top of the leaf. The bottom scraping shows similar results to the top, and similar to directly onto the leaf, however scraping showed the best results. Analysis was also conducted on the Town euc no rain and lambs ear (*Stachys byzantina*) with very hairy leaves located next to the town euc, and the results are similar to those shown here.

## Notes

### Competing Interest Statement

The authors have declared no competing interest.

### Summary of Updates

Minor editorial and typographical errors amended.

